# AD-HIES patients retain preTh17-cells that produce IL10 with opportunistic pathogens and induce IgE

**DOI:** 10.1101/2024.08.01.606136

**Authors:** Giorgia Moschetti, Chiara Vasco, Francesca Clemente, Paola Larghi, Sara Maioli, Edoardo Scarpa, Elena Carelli, Nadia Pulvirenti, Maria Lucia Sarnicola, Mariacristina Crosti, Manal Bel Imam, Willem van de Veen, Loris Rizzello, Sergio Abrignani, Lucia A. Baselli, Rosa Maria Dellepiane, Maria Carrabba, Giovanna Fabio, Jens Geginat

## Abstract

**Background:** Autosomal Dominant-Hyper-IgE Syndrome (AD-HIES) is caused by dominant-negative (DN) STAT3 mutations and characterized by high IgE levels, a lack of Th17-cells and recurrent infections with extracellular pathogens. We previously identified an enigmatic population of IL-10 producing CCR6^+^B-helper T-cells and investigated here their relationship to Th17-cells and STAT3 signalling requirements.

**Methods:** Human blood lymphocytes were analysed by multiparametric flow cytometry in healthy donors and AD-HIES patients. Analysis was performed by conventional gating or with bioinformatic tools. FACS-purified T-cell subsets were activated *in vitro* and Th17 differentiation assessed. T-cell antigen specificities were assessed by activation with heat-killed pathogens or antigenic peptide pools. B helper capacities were determined according to antibody secretion in B-T co-cultures by ELISA.

**Results:** CCR6^+^Th-cells that lacked subset-defining differentiation markers were mostly non-polarised central memory T-cells (T_CM_) that produced IL-10 and expressed RORγt. They were pre-committed to a Th17 fate, since TCR stimulation in the absence of polarising cytokines induced efficient Th17 differentiation. The latter was promoted by an autocrine loop of STAT3-activating cytokines. CCR6^+^Th-cells were reduced in patients with DN-STAT3 mutations but contained activated CCR6^+^T_CM_ that produced IL-10 and responded vigorously to AD-HIES-associated pathogens. These residual CCR6^+^Th-cells provided B-cell help for IgG and IgE production.

**Conclusions:** Th17 differentiation in AD-HIES patients was not completely impaired but arrested at an intermediate stage of IL-10 producing “pre-Th17”-cells. Surprisingly, DN-STAT3 mutations did not inhibit IL-10 production by CD4^+^T-cells. Pre-Th17-cells were activated by AD-HIES-associated pathogens and possessed B-helper functions, suggesting that they are not protective but promote aberrant IgE production.

## INTRODUCTION

CD4^+^ memory T-cells can be subdivided into two broad subsets of central and effector memory cells (T_CM_ /T_EM_)^1^. T_CM_ possess low effector functions, express CCR7 and CD62L that mediate homing to secondary lymphoid organs, and the stemness-promoting transcription factors TCF7/LEF1. Conversely, T_EM_ produce high levels of effector cytokines and express pro-inflammatory chemokine receptors that enables them to home to inflamed non-lymphoid organs. We previously identified human T_CM_ subsets that differentiate preferentially to Th1- and Th2-effector cells, termed pre-Th1 and pre-Th2-cells^2^. Later, Th17-cells and Tfh-cells have been identified that fight extracellular bacteria and fungi and promote B-cell antibody responses, respectively. Tfh express the chemokine receptor CXCR5 and home to B-cell follicles to provide help^3^. Th17-cells express the transcription factor ROR-γt, the chemokine receptor CCR6, and produce IL-17, IL-22^4,5^ and GM-CSF^6^. Th17-cells are highly heterogeneous^7–10^. In humans, conventional Th17-cells can be tacked according to CCR6 and CD161 co-expression^11^, and Th17 effector cells express in addition CCR5^12^. CCR6^+^Th-cells co-expressing CXCR3 produce IFN-γ and IL-17 and are called Th1/17-cells^13–15^. CCR6^+^T-cells co-expressing CCR10 produce IL-22, but not IL-17, suggesting that they represent Th22-cells^16,17^. Finally, CCR6^+^Th-cells in human blood co-expressing CXCR5 possess B-helper functions and are called Tfh17-cells^18^. Both Tfh-cells and Th17-cell differentiation is promoted by cytokines that signal via STAT3, like IL-6^19^ and IL-21^20,21^. The *in vivo* relevance of STAT3 in human T-cell differentiation is evident in patients with autosomal dominant Hyper-IgE Syndrome (AD-HIES), which contain dominant negative (DN) mutations of STAT3^22,23^. Deficient STAT3 signalling and its broad expression explain the multisystem manifestations of AD-HIES syndrome including dermatitis, pulmonary disease, vasculopathy, and skeletal and connective tissue abnormalities. High levels of serum IgE, eczema and recurrent infections are the hallmarks of AD-HIES. Severe bacterial infection, cold abscess, osteomyelitis, chronic muco-candidiasis and invasive fungal infections are common^24^. The recurrent infections in AD-HIES are explained by the lack of IL-17 producing T-cells^25,26^. We previously identified a Th-cell subset that expressed CCR6, but produced IL-10 rather than IL-17^27^ and possessed B helper functions^28^. Here we show that this enigmatic T-cells are T_CM_ which are pre-committed to a Th17 fate (pre-Th17). Surprisingly, they are generated in AD-HIES patients, are activated by AD-HIES-associated pathogens to produce IL-10 and maintain B-helper functions.

## Material and Methods

### Patients

AD-HIES was diagnosed according to the 2014 protocol by the Italian Primary Immunodeficiency Network (IPINet) group of the Italian Association of Pediatric Haematology Oncology (AIEOP) and patients of this study are part of the Italian AD-HIES cohort^29^. All patients carried a pathogenic DN mutation of STAT3. Genetics and clinical features of patients are shown in Figure 3A. All patients are followed according to the IPINet protocol. The local ethics committee approved this study (1097_2019bis, protocol ID #1305) and informed consent was obtained from all patients. This study adhered to the ethical principles outlined in the Declaration of Helsinki and followed the guidelines of Good Clinical Practice.

### Flow Cytometry

Human peripheral blood mononuclear cells (PBMCs) were isolated by standard Ficoll-Hypaque density gradient centrifugation. PBMC were stained with a combination of fluorochrome-conjugated antibodies as specified in sTable 1. Acquisition of stained cells was performed with a FACSymphony™ (Becton Dickinson, BD) and data analyzed with the FlowJo software (BD, version 10.8.1). To assess cytokine producing capacities, CD4^+^T-cell subsets were sorted on a FACSAria III (BD) and stimulated with 50ng/ml phorbol 12-myristate 13-acetate (PMA) and 500 ng/ml ionomycin for 4 hours in the presence of 10μg/mL Brefeldin A (all from Sigma Aldrich) in the last 2 hours. Cells were incubated with the Intracellular Fixation & Permeabilization Buffer Set (Thermo Fisher) and stained for intracellular cytokines (sTable 1). IL-10 production was also assessed in FACS-purified T-cell subsets following 30 hours stimulation with immobilized anti-CD3 antibodies (5µg/ml; Biolegend) and 10ng/mL IL-2 (Miltenyi) with an IL-10 secretion assay (Miltenyi), and after 18 hours of stimulation of PBMC with Staphylococcal enterotoxin B (SEB). Cytokine receptor expression was assessed following stimulation with 10ng/ml IL-6 or IL-23 (Miltenyi) at 37°C for 30 minutes. Cells were stained with anti phospho-STAT antibodies to assess STAT phosphorylation^20,30^.

### Immunofluorescence and imaging

IL-10 expression was analyzed in co-cultures of sorted CCR6^+^T-cell subsets, autologous monocytes and SEB. Experimental details of immunofluorescence are provided in the supplementary Material and Methods section. Fluorescence images were acquired using an inverted STELLARIS 8 Confocal Microscope (Leica Microsystems) equipped with a 405 laser and a tunable pulsed white light laser, utilizing a 40x air objective (numerical aperture [NA] 0.95) in three independent experiments. Quantification was performed using NIS-Elements V.5.30 software (Lim-Instruments). Ad-hoc designed pipelines of digital imaging analysis were implemented using the General Analysis 3 module of NIS-Elements to determine the intensity of fluorescence associated to the single cells present in the images.

### Th17 differentiation

CCR6^+^ and CCR6^-^T-cell subsets were sorted according to surface receptors as reported^28^. To obtain non-polarized cells, T-cell subsets were stimulated with PMA and Ionomycin, a cytokine secretion assay performed and IL-17A^-^IFN-γ^-^ or IL-17^-^IL-22^-^ cells sorted. Cells were stimulated with anti-CD3/28 beads (Invitrogen) at a 1:1 ratio. Recombinant cytokines (TGF-β, IL-4, IL-12 (R&D) or WNT10a, (Peprotech)) were added at 10ng/ml, antagonists of cytokine receptors at 10μg/ml (anti-IL-10, anti-IL-6 (Biolegend.) or 1μg/ml (anti-IL-10R and sIL-21R (R&D). After 6 days, cells were re-stimulated with PMA and ionomycin and analysed by cytokine staining. IL-6 production was measured in 6d supernatants of anti-CD3/28-stimulated T-cells by ELISA assay (Tebu Bio).

### Bioinformatic Analysis

Flow cytometry data were imported into FlowJo software (version 10.8.0) and a random down-sample of 6000 events for CD4^+^ T-cells was performed. The FCS file was analyzed by the R software. Sample batches were read using read.flowSet (2.6.0) from the flowCore R package. We applied the Logicle transformation that allows multiple samples to estimate transformation parameters. To reduce batch effects we normalized the signal of each marker with the function gaussNorm from the flowStat package (4.6.0). Samples were concatenated into a SingleCellExperiment object in R using the function prepData from the CATALYST R package. We performed high-resolution, unsupervised clustering and meta-clustering using FlowSOM (2.2.0) and ConsensusClusterPlus (1.58.0) package^31^. Manually annotated clusters were subsequently visualized on UMAPs. Functional pseudo-time analysis to infer the differentiation trajectory of cells was carried out by Diffusion Maps using the function run Diffusion Map on the SingleCellExperiment object using default parameters.

### Antigen specificity assay

Stimulation with heat-killed pathogens or pathogen-derived peptide pools was performed for 3 hours with 10^6^ PBMCs in RPMI 1640 complete medium supplemented with 5% human serum. Monensin (BD GolgiStop™, BD Biosciences) was added for an additional 15 hours of incubation and cells were analyzed for the co-expression of CD69 and cytokines. Heat-killed pathogens and antigenic peptide pools are listed in sTable 2.

### B helper assay

Naïve, CCR6^+^ and CXCR5^+^CD4^+^T-cells were isolated from total PBMCs of AD-HIES patients or healthy donors by FACS sorting (FACSAria III, BD). T-cells were co-cultured in V-shaped 96 wells plates with autologous CD20^+^B-cells at a 1:1 ratio in RPMI supplemented with 10% serum (Euroclone), 200U/ml IL-2 and Staphylococcal Enterotoxin B (SEB 100 ng/ml Merck). At day 14, co-cultures were supplemented with fresh medium and supernatants were harvested at day 14. Supernatants were tested for total IgG production by ELISA (Biolegend, Thermofisher Scientific, Abcam). To analyse IgE production, B-T co-cultures were performed with CXCR3^-^B helper subsets at a 3:1 B-T ratio with 1μg/ml SEB in the presence of 200 U/ml IL-2 and 10 ng/ml IL-4. Supernatants were tested by sandwich ELISAs for the secretion of IgG (BD) and IgE (Capture antibody: BD; Detection antibody: Invitrogen; Standard: Abcam).

### Statistical analysis

Statistics were performed with the Prism software (version 7; GraphPad Software, Inc., La Jolla, CA, USA). Statistical significance was evaluated using a Student t-test for the comparison of two groups with Mann-Whitney’s or Welch’s correction to analyze variables that were not normally distributed or by One way-ANOVA for comparison of more than two groups (Tukey’s post hoc correction). Significances were reported as *p< 0.05, **p< 0.01, ***p< 0.001. Error bars report +/-SEM in all Figures.

## RESULTS

### CCR6^SP^T-cells are central memory T-cells (T_CM_) that express ROR-γt and produce IL-10

We previously identified an enigmatic population of CCR6^+^ B-helper T-cells cells that lacked gene signatures and established surface markers of Th17-, Th1/17- and Tfh17-cells called CCR6^“single-positive”(SP)^T-cells^28^. To map the differentiation stage of CCR6^SP^T-cells, we first performed a deep phenotypic analysis in the blood of healthy donors (Figure 1A, sFigure 1A). CCR6^SP^T-cells expressed low levels of the gut-homing receptors CCR9 and β7-integrin and lacked CXCR6. They expressed however several receptors involved in skin-homing, including CCR10, CLA, β1-integrin and CCR4 (Figure 1A, sFigure 1B). Since human CCR6^+^CCR10^+^T-cells contain Th22-cells^7,16,17^ we analysed them separately and focussed the further analysis on CCR10^-^CCR6^SP^-T-cells (sFigure 1A). Most CCR6^SP^T-cells displayed a T_CM_ phenotype, since they co-expressed CCR7 and CD62L (Figure 1B). Moreover, they expressed high levels of CD27 and lacked PD1 (sFigure 1C). TCF7 and LEF1 are transcription factors of the WNT pathway that promote stemness. CCR6^SP^T-cells expressed intermediate levels of TCF7/LEF1 (Figure 1C), as is characteristic for T_CM_ (sFigure 1D). Expression of ROR-γt, the lineage-defining transcription factor of Th17-cells, was restricted to CCR6^+^T_CM_ and T_EM_ (sFigure 1E). Among CCR6^+^T-cells it was expressed at high level in Th17-cells and Th1/17-cells, but it was also expressed in CCR6^SP^T-cells (Figure 1D). We then assessed cytokine producing capacities of CCR6^+^T-cell subsets upon *ex vivo* stimulation with PMA and Ionomycin (Figure 1E). Th17-, Th1/17- and Th22-cells produced as expected preferentially IL-17, IFN-γ and IL-22, respectively^13,16,18^. Th1/17- and Th22-cells produced also high levels of GM-CSF. CCR6^SP^T-cells and Tfh17-cells produced only low levels of effector cytokines, suggesting that they were predominantly non-polarized. Since PMA plus Ionomycin stimulation is suboptimal to induce IL-10 production in human CCR6^+^Th-cells^27^, we stimulated CCR6^+^T-cell subsets with monocytes and SEB and analysed IL-10 production by immunofluorescence (Figure 1F, sFigure 1F). IL-10 was produced at significantly higher levels by CCR6^SP^-T-cells as compared to Tfh17-cells, and Th17-cells produced only low levels. Finally, since IL-6R and IL-23R promote respectively early and late Th17 differentiation^12,19,32,33^ we assessed their expression according to ligand-induced STAT phosphorylation^20^ (Figure 1G, sFigure 1G). IL-6 induced STAT phosphorylation in most CCR6^SP^T-cells, whereas IL-23 largely failed to do so.

**FIGURE 1:**
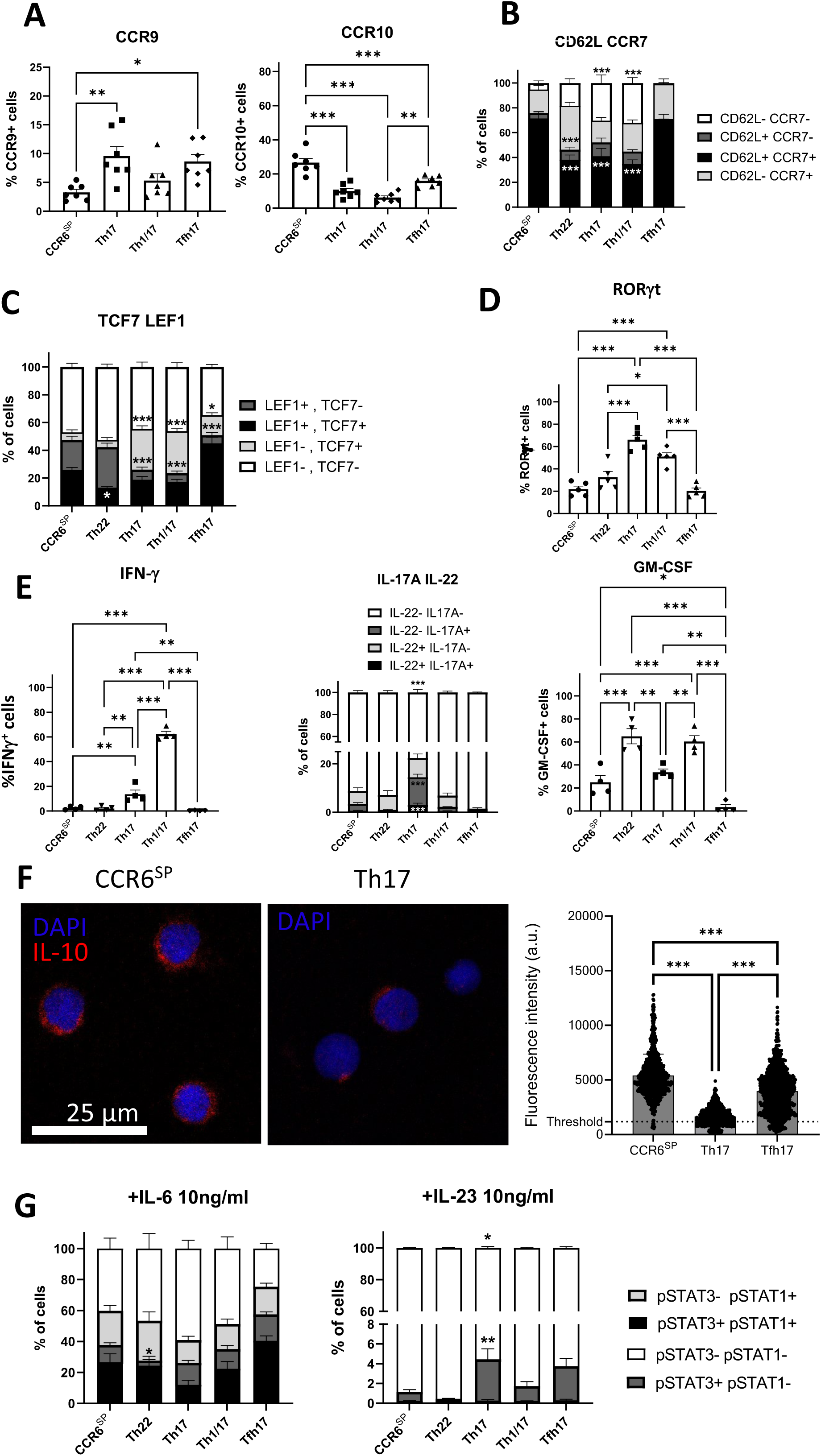
CCR6^SP^-T-cells are T_CM_ that express ROR-γt and produce IL-10. **A:** *Ex vivo* frequencies of CCR9- or CCR10-expressing cells among gated CD4^+^CCR6^+^Th-cell subsets (see gating strategy in sFigure 1A) in healthy donors (n=7) Expression of **B:** CD62L and/or CCR7 (stacked histograms, n=7; Th22: CCR6^+^CCR10^+^). **C:** TCF7 and/or LEF1 (n=7) and **D:** RORC/γt in healthy donors (n=5) **E:** Production of IFN-γ, IL-17 and/or IL-22 and GM-CSF following 4 hour stimulation of the indicated FACS-purified CCR6^+^T-cell subsets from healthy donors with PMA and Ionomycin (n=4) **F:** IL-10 production by the indicated CCR6^+^T-cell subsets from healthy donors upon stimulation with monocytes and SEB for 16 hours was analysed by immunofluorescence. At least 1600 cells from 3 different ì donors were analysed. **G:** Expression of functional cytokine receptors in T-cell subsets from healthy donors was assessed after stimulation for 30 minutes with the relevant cytokines by STAT phosphorylation. Statistical significant differences were calculated by One-way ANOVA and reported as *p< 0.05, **p< 0.01, ***p< 0.001. */*** on the top of stacked histograms report significant differences of double-negative cells.

### An autocrine loop of STAT3-activating cytokines promotes TCR-induced Th17 differentiation of CCR6^SP^T-cells in the absence of polarising cytokines

Since CCR6^SP^T-cells expressed ROR-γt we investigated if they were pre-committed to a Th17 fate. To exclude contributions of contaminating effector T-cells, we purified non-polarized T-cells by depleting cells that could produce IL-17, IFN-γ or IL-22 *ex vivo* with a cytokine secretion assay. The capacity to produce the depleted cytokines was then analysed after *in vitro* culture in the absence of Th17-polarizing cytokines for 7 days. CCR6^SP^T-cells acquired high levels of IL-17 and IL-22 producing capacities under these neutral conditions, whereas CCR6^-^ control Th-cells did not (Figure 2A). Notably, CCR6^SP^T-cell produced also significantly higher amounts of IL-17 as compared to Tfh17 and Th1/17-cells (Figure 2B). To analyse the requirements of this spontaneous Th17 differentiation, we added cytokines that promote alternative T-cell fates. Acquisition of IL-17 and IL-22 producing capacities by CCR6^SP^T-cells was weakly inhibited by TGF-β and WNT10A (Figure 2C and sFigure 2A), which activates LEF1/TCF7^34^. Conversely, IL-4 strongly inhibited Th17 differentiation and IFN-γ production. IL-12 had a weak inhibitory effect on IL-17 production but strongly induced IFN-γ. Consequently, it promoted co-production of IFN-γ and IL-17, as is characteristic for Th1/17-cells^13,20,35^ (Figure 2C, sFigure 2B). Since Th17 differentiation requires STAT3, we investigated the role of cytokines that are produced by TCR-activated CCR6^SP^T-cells and activate STAT3, such as IL-6 (sFigure 2C), IL-10 (Figure 1F^28^) and IL-21^20^. TCR-activated CCR6^SP^T-cells contained indeed high levels of phosphorylated STAT3 in the absence of exogenous cytokines (Figure 2D). Moreover, STAT3 phosphorylation was significantly inhibited when IL-6, IL-10 and IL-21 were neutralized (Figure 2D). Importantly, neutralization of these STAT3-activating cytokines also significantly reduced IL-17 and IL-22 production (Figure 2D, sFigure 2D).

**FIGURE 2:**
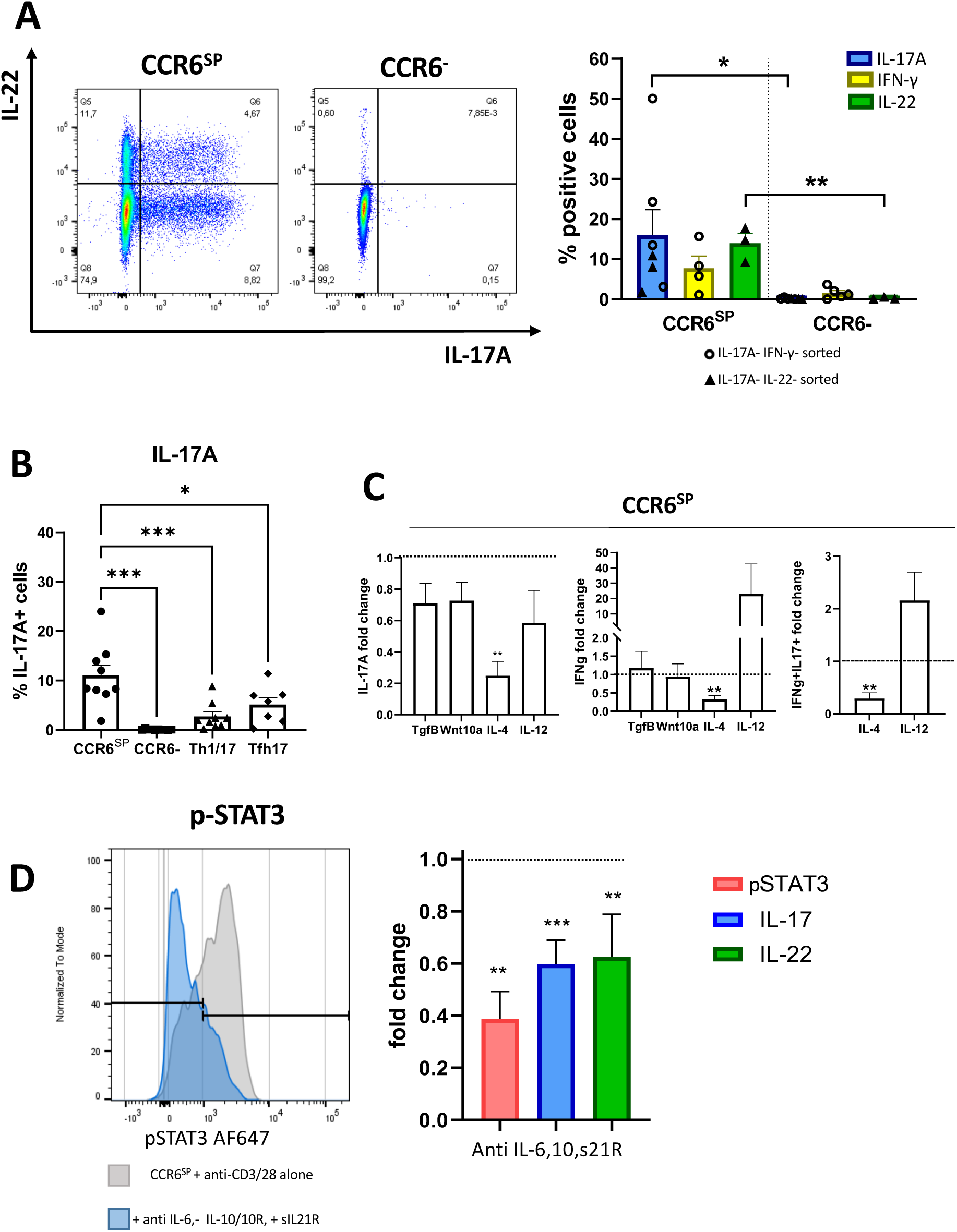
TCR-activated CCR6^SP^T-cells differentiate spontaneously to Th17-cells. **A.** FACS-purified CCR6^SP^ T-cells or CCR6^-^ control cells from healthy donors were depleted from residual IL-17 and IFN-γ (n=4) or IL-17 and IL-22 producing cells (n=3) with a cytokine secretion assay. Cells were activated with anti-CD3/28-coated beads and the induction of the previously depleted cytokines by PMA plus Ionomycin was analyzed by intracellular staining. Left: Dot plots of IL-17 and IL-22 induction in one representative donor. Right: Bars report the frequencies of cytokine-producing cells. IL-17^+^ and IFN-γ^+^-depleted cells are indicated by filled circles, IL-17^+^ and IL-22^+^ depleted cells by triangles. Statistical significance for each cytokine was calculated with a paired student’s t-test. **B.** IL-17 production by the indicated T-cell subsets cultured for 6 day with anti-CD3/28-coated beads induced by PMA and Ionomycin. Statistical significance was calculated with One-way ANOVA. **C.** FACS-purified total CCR6^SP^T-cells from healthy donors were cultured for 6 day with anti-CD3/28-coated beads in the absence or presence of the indicated agonists and were analyzed for IL-17 and IFN-γ producing capacities. **D:** FACS-purified CCR6^SP^T-cells from healthy donors were cultured with anti-CD3/28-coated beads in the absence or presence of antagonists of IL-6, IL-10 and IL-21 and analyzed for STAT3 phosphorylation after 24 hours or for IL-17 and IL-22 production after 6 days following brief restimulation with PMA and Ionomycin. **C/D:** Shown is the fold-change as compared to CCR6^SP^T-cells stimulated with anti-CD3/28 beads alone (indicated by dotted lines). Statistical significance for each condition was calculated with a paired student’s t-test and is indicated by */**/***/**** on the top of the columns.

### Residual CCR6^+^T-cells in AD-HIES contain Th1/17- and CCR6^SP^T-cells and produce IL-10

To address the role of STAT3 in CCR6^SP^T-cells *in vivo*, we then analysed CD4^+^T-cells from AD-HIES patients (Figure 3A)^36^. CD4^+^T-cells from AD-HES patients were compared to healthy donors (HDs) by multi-dimensional flow cytometry. Unsupervised principal component analysis (PCA) revealed significant differences (Figure 3B). Unified Manifold Approximation and Projection (UMAP, Figure 3C, sFigure 3A-C) identified six CCR6^+^Th-cell clusters, which were reduced in AD-HIES. Notably, cluster 9 contained CCR6^SP^T-cells and co-clustered with Th1/17_CM_ (Cluster 7) and Tfh (Cluster 8), but not with CCR6^+^CCR10^+^ clusters (Clusters 11 and 13, sFigure 3A). The reduction of CCR6^+^Th clusters was compensated by an increase of naïve T-cells (Cluster 1, Figure 3D), whereas Th1 clusters (Clusters 6, 16) were unaffected. Diffusion map analysis suggested a progressive differentiation from naïve to effector T-cells, with T_CM_ and T_EM_ being respectively early and late differentiation intermediates (Figure 3E). Moreover, the diffusion map was consistent with an early-to-intermediate differentiation stage of CCR6^SP^T-cells, positioning them between T_FH_- and other T_CM_ clusters. Manual gating confirmed that AD-HIES patients exhibit an increase of naïve T-cells and a reduction of all CCR6^+^T-cell subsets (sFigure 4A). Conversely, the frequencies of Th1, Th2 and of regulatory T-cell subsets, i.e. FOXP3^+^Tregs and EOMES^+^Tr1-like cells^37^, were similar (sFigure 4B). Notably, while Th17- and Tfh17 subsets were largely undetectable in AD-HIES, Th1/17-cells and CCR6^SP^ were only partially reduced (sFigure 4A). Consequently, when the relative contributions of subsets to CXCR3^-^ CCR6^+^Th-cells were analysed, most cells in AD-HIES patients displayed a CCR6^SP^ phenotype (Figure 4A). Consistently, CCR6^+^T-cells from AD-HIES patients contained higher frequencies of T_CM_ (Figure 4B), expressed higher levels of LEF1/TCF7 (Figure 4C) and lower levels of ROR-γt (Figure 4D). Notably, CCR6^+^T-cells from AD-HIES patients expressed in mean higher levels of CCR9 and significant lower levels of β1-Integrin (Figure 4E). The expression of other homing receptors was not significantly different (sFigure 4C). Finally, CCR6^+^T-cells in AD-HIES lacked IL-17 producing capacities, whereas production of IL-21 (Figure 4F), IFN-γ, IL-2, IL-4, TNF and GM-CSF were similar (sFigure 4D). They produced however significantly higher levels of IL-10 (Figure 4F). This was also true upon more physiological TCR stimulation with anti-CD3 antibodies^27^ or with the superantigen SEB^38^ (sFigure 4E). Conversely, IL-10 production by CCR6^-^Th-cells (Figure 4F) and by regulatory T-cells^37^ was not significantly different (sFigure 4F).

**FIGURE 3:**
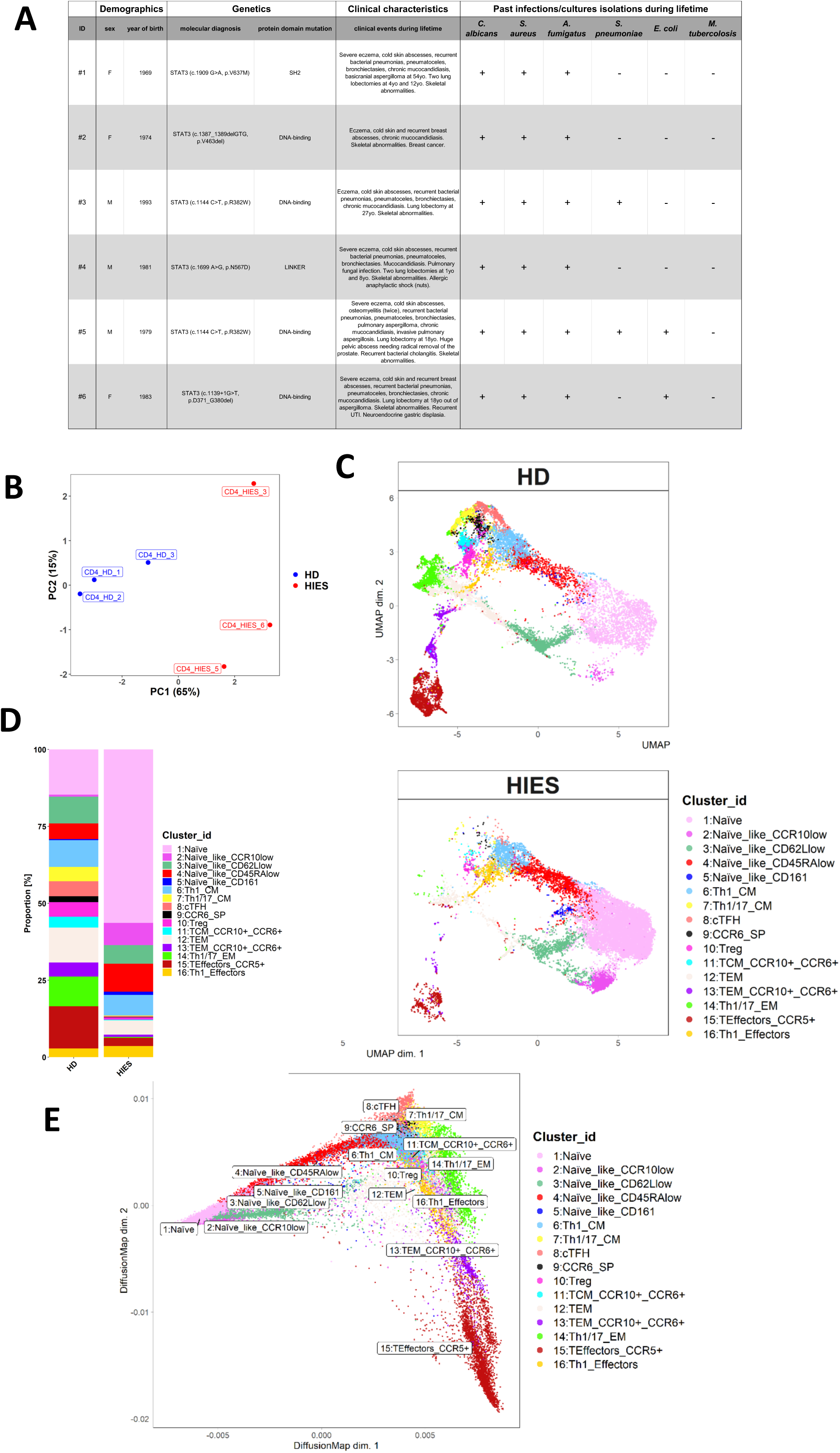
Unbiased analysis of the CD4^+^T-cell compartments unveils a selective decrease of CCR6^+^T-cell clusters in AD-HIES. **A** Table showing mutations and clinical features of HIES patients **B.** Principal Component Analysis of 3 AD-HIES patients and 3 healthy donors (HDs), respectively coloured in blue and in red. Number in the label indicated the sample IDs. **C.** UMAP maps of the 16 identified CD4^+^T-cell clusters marked with different colours in the blood of HD and AD-HIES patients. **D.** Bar-plots showing cluster distributions in the blood of HD and AD-HIES patients. **E.** Diffusion Map analysis of predicted cell differentiation pathways of the 16 identified CD4 T-cell clusters in pseudotime**. C-E** Colour code and names of the 16 clusters are indicated in the Figure.

**FIGURE 4:**
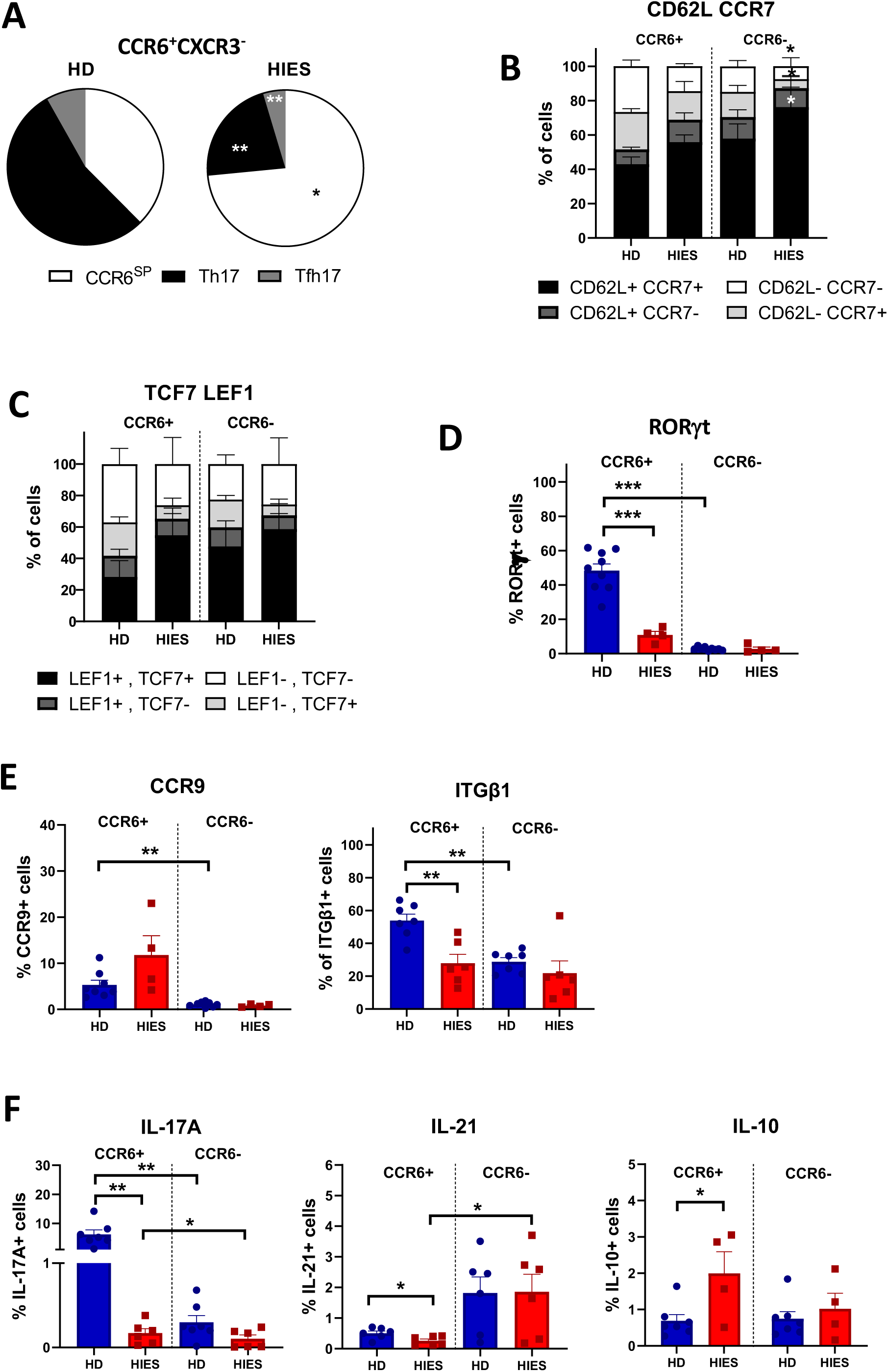
Residual CCR6^+^T-cells cells in AD-HIES patients display features of CCR6^SP^T-cells and produce increased amounts of IL-10. **A:** Pie charts show the relative contributions of phenotypical Tfh17-cells, Th17-cells and CCR6^SP^T-cells to CXCR3^-^CCR6^+^T-cells. **B-E:** Gated CCR6^+^ and CCR6^-^CD4^+^T-cells in HD (n=5-9) and AD-HIES patients (n=4-6) were analyzed *ex vivo* by flow cytometry for the expression of **B.** CD62L and/or CCR7 **C.** LEF1 and/or TCF7 **D:** RORγt and **E:** CCR9 and β1-integrin. **F:** Frequencies of IL-17A, IL-21 and IL-10 expressing cells among CCR6^+^ and CCR6^-^T-cells stimulated *ex vivo* with PMA and Ionomycin obtained from AD-HIES patients (red bars, n= 4-6) and HD (blue bars, n=7). **B-E:** Statistical significance was calculated between HDs and HIES patients and between CCR6^+^ and CCR6^-^ subsets of the same individuals using unpaired or paired t-tests, respectively.

### CCR6^+^T-cells in AD-HIES are activated by opportunistic pathogens to produce IL-10

CCR6^+^T-cells from AD-HIES patients were activated *in vivo*, as indicated by increased expression of PD1, CD69 and of the proliferation marker Ki67 (Figure 5A). We investigated if they might be activated by AD-HIES-associated infections. Notably, all analysed patients had recurrent infections with Staphylococcus aureus, Candida albicans and Aspergillus fumagatus (Figure 3A). Moreover, some patients experienced infections with Escherichia coli and/or Streptococcus pneumoniae, whereas infections with Mycobacterium tuberculosis were never diagnosed. We could detect IL-17 production by CCR6^+^T-cells in response to all these heat-killed pathogens in healthy donors, but not in AD-HIES patients (Figure 5B). However, CCR6^+^T-cells from AD-HIES patients produced increased amounts of IL-10 that reached statistical significance for most pathogens. CCR6^+^T-cells from AD-HIES patients produced also significantly more IL-2 and IFN-γ in response to Escherichia coli^12^, Streptococcus pneumoniae and Mycobacterium tuberculosis^13^ (Figure 5B). Conversely, CCR6^-^ control T-cells responded poorly (sFigure 5A). To control for possible bystander T-cell activation by heat-killed pathogens, we stimulated with LPS. However, LPS failed to induce T-cell cytokine production (sFigure 5B). As an additional control, we analysed CD4^+^T-cell responses to antigenic peptides derived from Candida albicans (MP65), Aspergillus (SHMT) and from EBV. Consistent with the results obtained with heat-killed Candida, we observed a switch from IL-17 to IL-10 production in AD-HIES (Figure 5C, sFigure 5C). CCR6^-^T-cells responses were inconsistent (sFigure 5D). However, CCR6^-^T-cells produced some IFN-γ in response to EBV-derived peptides in HDs and AD-HIES patients (sFigure 5E), indicating that virus-specific Th1-cells were intact^14^.

**FIGURE 5:**
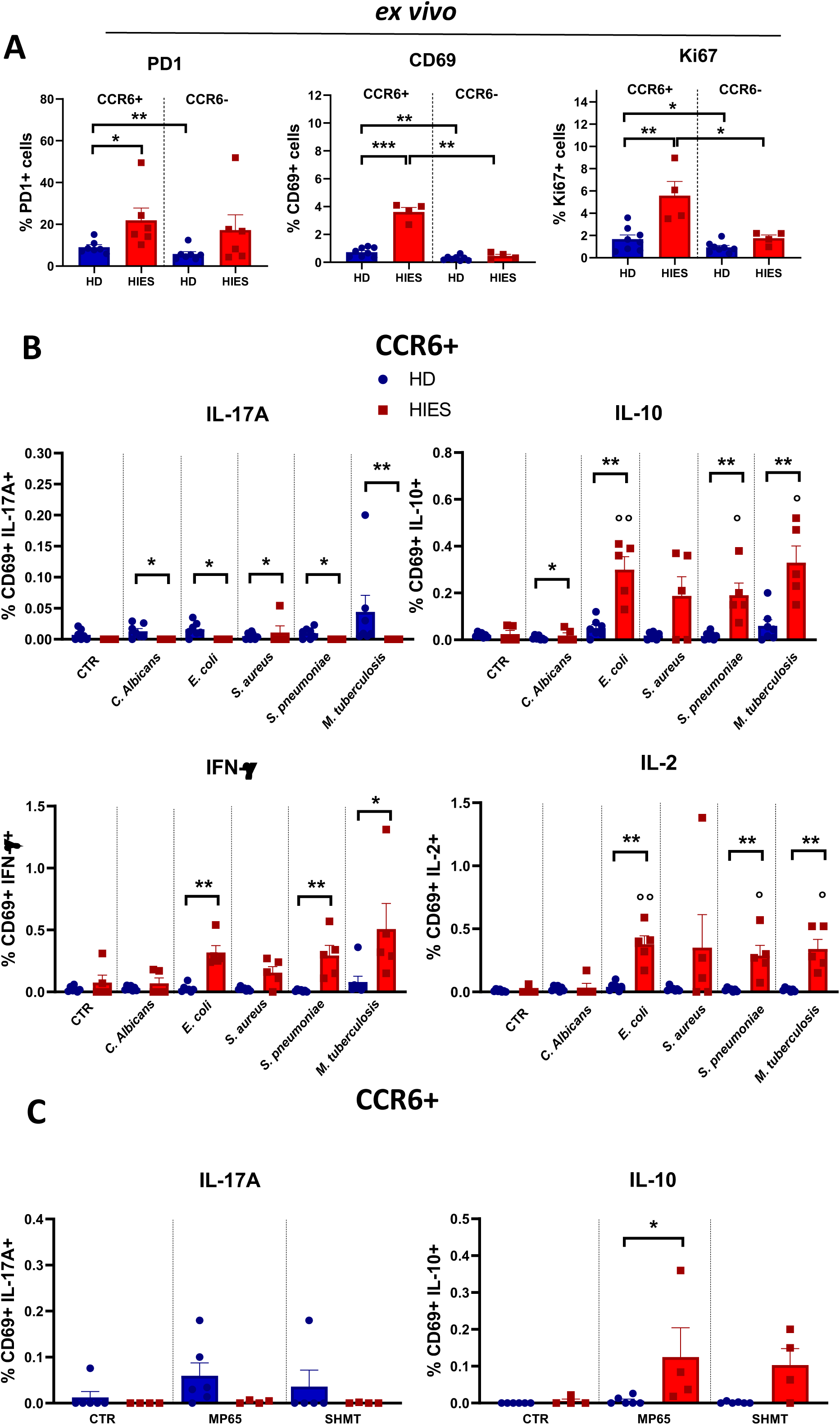
CCR6^+^T-cells in AD-HIES patients are activated *in vivo* and produce IL-10 in response to opportunistic pathogens. **A:** *Ex vivo* expression of PD1 (left), CD69, (center) and Ki67 (right) in gated CCR6^+^ and CCR6^-^T-cells from AD-HIES patients (red bars, n= 6) and HD (blue bars, n=7). **B.** Co-expression of CD69 and IL-17A (upper left panel), IL-10 (upper right panel), IFN-γ (lower left panel) and IL-2 (lower right panel) in CD4^+^CCR6 T-cells from AD-HIES patients (red bars, n=5) and HD (blue bars, n=8) stimulated with the indicated heat-killed pathogens. Cells incubated without pathogens were analysed as negative control (CTR). **C.** Co-expression of CD69 and IL-17A or IL-10 in CCR6^+^T-cells from AD-HIES patients (red bars, n=4) and HD (blue bars, n=6) induced by antigenic peptide pools derived from Candida albicans (MP65) or from Aspergillus fumagatus (AMT). Cells incubated without peptides were analysed as negative control (CTR). Statistical significances were calculated with student’s t-test.

### CCR6^+^T-cells in AD-HIES possess B-helper functions and promote IgE production

CCR6^+^Th-cells possess B-helper functions in STAT3-sufficient individuals^28^ and we investigated here if they could induce antibody production also in patients with DN-STAT3 mutations. Their B-cell compartments was characterized by a strong decrease of memory cells^39^, which was again compensated by an increase of naïve cells (Figure 6A, sFigure 6A). However, they contained higher frequencies of plasmablasts, suggesting ongoing B-cell activation (Figure 6B, sFigure 6B). We co-cultured autologous B- and Th-cells in the presence of SEB and measured antibody production by ELISA. CCR6^+^Th-cells from AD-HIES patients induced significant amounts of IgG, demonstrating that they possessed B-helper functions (Figure 6C). Notably, CXCR5^+^Tfh-cells possessed comparable B-helper capacities, whereas naïve CD4^+^T-cells were inefficient. IgE production was undetectable under these conditions. We therefore performed optimised T-B co-cultures in the presence of IL-4. We also excluded CXCR3^+^T-cells, because they lack B helper functions^18^ and because we observed increased B-helper activities by CXCR3-depleted Th-cells in preliminary experiments (sFigure 6C)^40^. IgE were induced by CXCR3^-^CCR6^+^T-cells both in healthy donors and in AD-HIES patients under these conditions (Figure 6D). Notably, IgE secretion induced by CXCR3^-^ CCR6^+^T-cells was similar to that induced by Tfh2-cells, which were analysed as positive control^18^. IgE levels were in mean lower in cultures of AD-HIES patients but this difference was not statistically significant. CCR6^+^T-cell-induced IgG levels were in contrast strongly and significantly reduced in AD-HIES (Figure 6D), consistent with a more prominent role of STAT3 for IgG induction.

**FIGURE 6:**
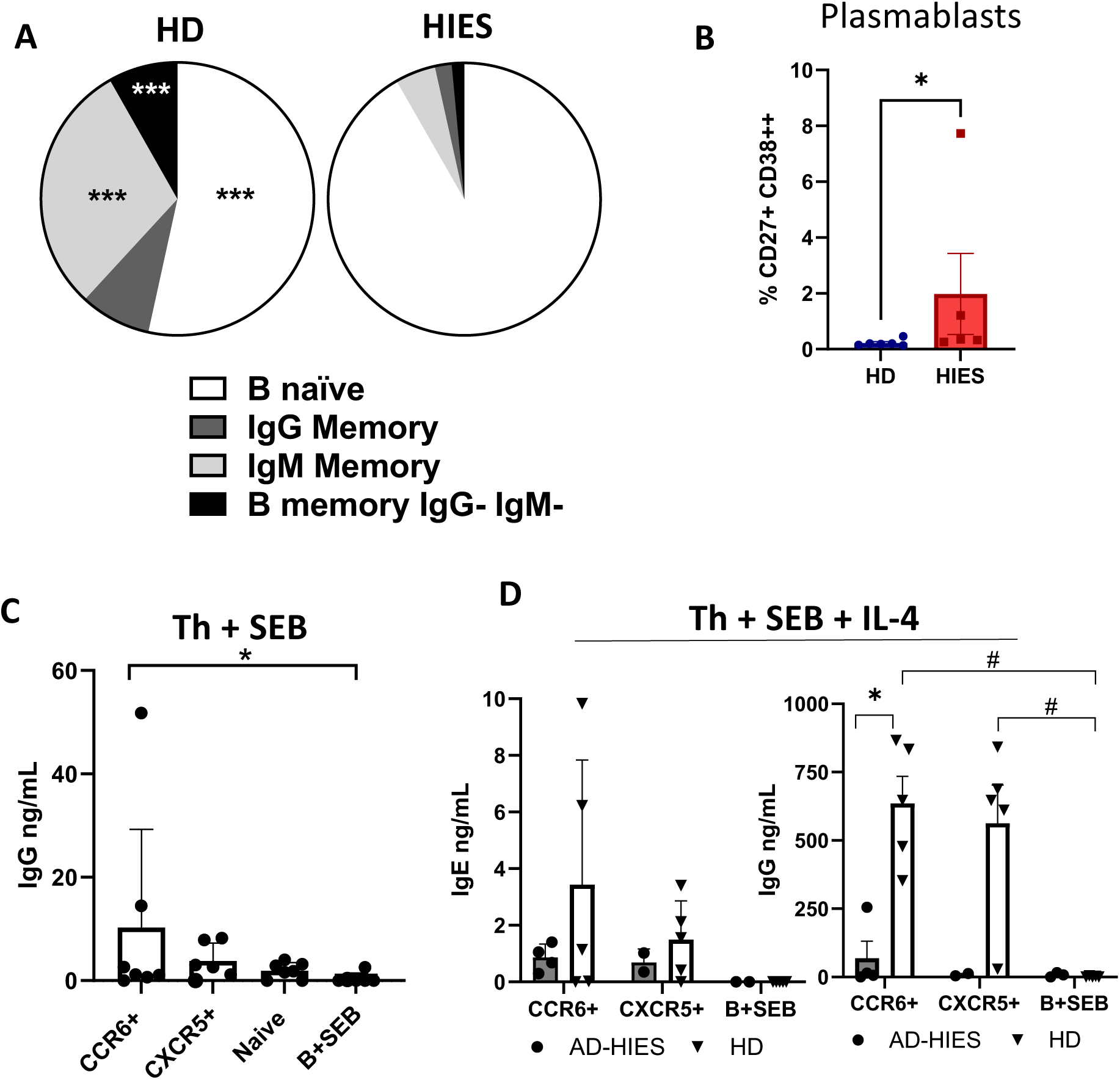
CCR6^+^T-cells from AD-HIES patients possess B helper capabilities and promote IgE production. **A.** Pie Charts show the compositions of the B cell compartments in healthy donors and AD-HIES patients. **B:** Frequencies of plasmablasts in HD and AD-HIES patients (CD19^+^CD27^hi^CD38^hi^). A/B: Statistical significances were calculated with an unpaired t-test. **C.** B-cells from AD-HIES patients were cultured with SEB in the absence or presence of the indicated FACS-purified Th-cells and IgG secretion was assessed by ELISA. **D.** B-cells from healthy donors and AD-HIES patients were cultured with SEB and IL-4 in the absence or presence of CCR6^+^CXCR3^-^Th-cells and Tfh2-cells and IgG and IgE secretion assessed by ELISA. C/D: Statistical significance between was calculated by a Man-Whitney Test.

## Discussion

We mapped here the differentiation stage of an enigmatic population of CCR6^+^B-helper T-cells that we identified previously^27,28^. These CCR6^+^Th-cells displayed a T_CM_ phenotype and expressed ROR-γt, indicating that they represent an early stage of Th17 differentiation. Consistently, they acquired IL-17 producing capacities upon TCR stimulation in the absence of exogenous Th17-promoting cytokines. This “spontaneous” Th17 differentiation was fast and highly efficient. Conversely, Th17 differentiation of uncommitted human helper T-cells is slow, inefficient and requires a combination of several Th17-promoting cytokines^30^. Thus, CCR6^SP^T-cells are pre-committed to a Th17 fate (“pre-Th17 cells”), in analogy to the pre-Th1 and pre-Th2 cells that we have identified previously^2^. A feature of these pre-committed T_CM_ is that they generate spontaneously effector cells of their respective lineage, but that they retain nevertheless a relevant degree of plasticity to differentiate to alternative fates upon instruction by polarising cytokines^41^. In the case of pre-Th17-cells, IL-12 induced IFN-γ, and promoted the generation of Th1/17-cells *in vitro*. Consistently, AD-HIES patients contained residual Th1/17-cells that produced IFN-γ in response to M. tuberculosis, explaining why M. tuberculosis is controlled in AD-HIES in contrast to ROR-γt-deficient patients^42^.

STAT3 signalling has a non-redundant role in Th17 differentiation, as demonstrated by the absence of IL-17 producing T-cells in humans and mice with genetic defects in STAT3 activation^23,25,36,43^. Naïve T-cells developing into Th17-cells activate STAT3 initially in response to IL-6, which then induces IL-21 production to amplify STAT3 activation and Th17 differentiation in an autocrine feed-forward loop^19,44,45^. Consistently, Th17 differentiation of pre-Th17-cells was inhibited when STAT3-activating cytokines were neutralized *in vitro* or when STAT3 signalling was impaired *in vivo,* i.e. in AD-HIES. Notably, TGF-β induces CCR6 and ROR-γt in human CD4^+^T-cells but inhibits IL-17 production^27,46^. Once induced, CCR6 expression in human CD4^+^T-cells is remarkably stable^47^. Thus, pre-Th17-cells may be generated with TGF-β, which also induces CCR4^48^ and IL-10^44^, in a at least partially STAT3-independent manner.

STAT3 signalling is also critical for Tfh differentiation. We observed a selective lack of Tfh17-cells in AD-HIES. Tfh1/2 subsets were present, indicating distinct differentiation requirements. Importantly, AD-HIES patients contained also residual CCR6^+^Th-cells, including pre-Th17-cells, and possessed B-helper functions. Interestingly, CCR6^+^Th-cells and Tfh2 induced similar levels of IgE in both STAT3-deficient and -sufficient individuals. It is thus tempting to speculate that CCR6^+^Th-cells contribute to IgE production not only in HIES, but also in allergy. Notably, they are however extrafollicular B-helper T-cells^28^ and may thus induce antibodies of lower quality, as is characteristic for AD-HIES^39^.

Inborn errors of immunity unveil non-redundant pathways for the immune defence against pathogens. AD-HIES patients lack Th17-cells and experience consequently recurrent infections with extracellular pathogens^26,49^. These opportunistic pathogens activated selectively the residual CCR6^+^T-cells to produce IL-10, but also IL-2 and IFN-γ. IL-10 producing Th17-cells specific for S. aureus have previously been described in healthy individuals^9^, but they lacked protective functions in mice^50^. Notably, since IL-10 activates STAT3 it is probably functionally irrelevant in AD-HIES. The increased IL-10 production by CCR6^+^T-cells in AD-HIES patients was nevertheless surprising, since IL-10 production in developing Th17-cells is promoted by IL-6 and IL-21 that signal via STAT3^41,44^. Moreover, IL-21 failed to induce IL-10 production by *in vitro* activated T-cells from AD-HIES patients^51^. Our results indicate however that STAT3 does not play a non-redundant role for T-cell IL-10 production *in vivo,* neither by helper nor by regulatory T-cells. Indeed, T-cell IL-10 production can also be induced by IL-4 or IL-12 via STAT4 or STAT6, respectively^52^, or by chronic TCR stimulation.

In conclusion, AD-HIES-associated pathogens induce the generation of pre-Th17-cells that produce IL-10. They cannot differentiate to Th17-cells in the absence of STAT3 signalling and consequently lack protective functions but may instead contribute to the aberrant IgE production.

## ACKNOWLEDMENTS

This work was supported by the Telethon Foundation, grant n. GGP19323 and by the Italian Ministry of Health PRIN 2022 Project P20228W4FJ. Both the Pediatrics and Adults Centres are involved in the European Reference Network for Rare Immunodeficiency, Autoinflammatory and Autoimmune Diseases, ERN-RITA, the study is part of the activities of the Centres. We thank AD-HIES patients and healthy donors that donated blood.

## Author contributions

G.M and C.V designed experiments, performed and supervised experiments, analysed the data and wrote the paper. F.C performed bioinformatic analysis. P.L. S.M, E. S., E. C., M. P, M.L.S, M.C, M. B. I. analysed and/or performed experiments. L.R. and S. A. provided critical Discussion and wrote the paper. L.A. B, R.M. D., M. C. and G. F. provided access to patients and their clinical characteristics and wrote the paper. J.G. designed the study, supervised experiments and wrote the paper.

<u>SUPPLEMENTARY FIGURE 1:</u>

**<u>A</u>**<u>: Gating strategy to identify CCR6^+^Th subsets in human blood</u> **B**: Frequencies of, Integrin-β7^+^, CXCR6^+^, CLA^+^, Integrin-β1^+^, CCR4^+^, CD27^+^ and PD1^+^ cells were measured *ex vivo* by surface staining on gated CCR6^+^T-cell subsets. **C**: Stacked histogram show co-expression of TCF7/LEF1 in CD4^+^ naïve (CD45RA^+^CCR7^+^CD95^-^), T_SCM_ (CD45RA^+^CCR7^+^CD95^+^), T_CM_ and T_EM_. **D.** RORγt expression in CCR6^+/-^ naïve (CCR7^+^CD45RA^+^), central (CCR7^+^CD45RA^-^, T_CM_) and effector memory (CCR7^-^CD45RA^-^, T_EM_) T-cells from healthy donors (n=8). Statistical significances were calculated using One way ANOVA. **E.** Immunofluorescence for IL-10 production in Th17-cells and Tfh17cells following stimulation with monocytes and SEB for 16 hours **F**: Negative controls for STAT1/3 activation in the absence of recombinant IL-6 and IL-23

<u>SUPPLEMENTARY FIGURE 2:</u>

**A**: FACS-purified CCR6^SP^-T-cells were stimulated with anti-CD3/CD28-coated beads in the absence or presence of recombinant TGF-β, WNT10A, IL-4 or IL-12 as indicated. After 6 days cells were restimulated with PMA and Ionomycin and production of IL-22 analysed. Statistical significance was calculated by One-way ANOVA **B**: FACS dot plots of one representative experiment showing IL-17 versus IFN-γ production by CCR6^SP^-T-cells stimulated with anti-CD3/CD28-coated beads in the absence or presence of IL-4 or IL-12. **C**: The day 6 culture supernatant of anti-CD3/CD28-stimulated CCR6^+^-T-cell subsets was analysed for IL-6 by ELISA (n=3). **D:** Dot plots of one representative experiment showing CCR6^SP^T-cells cultured with anti-CD3/28-coated beads in the absence or presence of anti-IL-10 antibodies and analyzed for IL-17 and IL-22 production.

<u>SUPPLEMENTARY FIGURE 3:</u>

**A**: Heatmap of median normalized expression of markers within each of the identified 16 clusters. Dendrogram on the left side of the heatmap shows similarities among clusters. Red indicates high and blue low expression. **B**: Histograms showing the expression levels of the 13 markers in the 16 CD4^+^T-cell clusters. **C:** UMAPs showing the expression of the individual 13 markers. Red indicates high, green intermediate and blue low expression levels.

<u>SUPPLEMENTARY FIGURE 4:</u>

**A:** Left: Frequencies of naïve (CD45RA^+^CCR7^+^CD95^-^) and memory T-cell subsets, i.e. Tscm (CD45RA^+^CCR7^+^CD95^+^), Tcm (CCR7^+^CD45RA^-^), Tem (CCR7^-^CD45RA^-^) and Temra (CCR7^-^CD45RA^+^) in HDs (blue bars) and HIES patients (red bars). Right: Frequencies of CCR6^SP^, Th17, Th1/17 and Tfh17 T-cells among CD4^+^T-cells in healthy donors (HD, blue bars, n=7) and AD-HIES patients (red bars, n=6). **B:** Frequencies of Th1-(CD4^+^CXCR3^+^CCR4^-^CCR6^-^) and Th2-cells (CD4^+^CXCR3^-^CCR4^+^CCR6^+^), as well as Tregs (CD4^+^FOXP3^+^IL-7R^lo^) and Tr1-cells (CD4^+^GZMK^+^IL-7R^lo^) in HD (blue bars, n=7) and AD-HIES patients (red bars, n=6). **C.** Expression of β7-integrin and CLA in gated CCR6^+^ and CCR6^-^Th-cells in HD and AD-HIES patients. **D**: Production of the indicated cytokines by *ex vivo* stimulated CCR6^+^ and CCR6^-^ T-cell subsets of healthy donors (HD, blue, n=7) and AD-HIES patients (red, n=4). **E:** Left: Frequencies of IL-10-producing cells after 30h stimulation with anti-CD3 antibodies and IL-2 of CD4^+^T-cell subsets sorted according to CCR6 expression from AD-HIES patients (red bars, n=4) and healthy donors (blue bars, n=6). Right: Frequencies of IL-10 producing cells among superantigen (SEB)-stimulated, gated CCR6^+^ and CCR6^-^T-cells from AD-HIES patients (red bars, n=5) and HD (blue bars, n=7). **F**: IL-10 production by *ex vivo*–stimulated T regulatory subsets from AD-HIES patients (red bars, n= 4-6) and healthy donors (blue bars, n=3-4). Statistical significances were calculated with t-tests.

<u>SUPPLEMENTARY FIGURE 5:</u>

**A**: Co-expression of CD69 and the indicated cytokines in CCR6^-^T-cells from AD-HIES patients (red bars, n=4-5) and healthy donors (HD, blue bars, n=7-8) upon stimulation with pathogen-derived antigens **B**: Co-expression of CD69 and the indicated cytokines in CCR6+ (upper panels) and CCR6^-^ (lower panels) T-cells from AD-HIES patients (red bars, n=4-5) and healthy donors (HD, blue bars, n=7-8) upon stimulation with LPS. Unstimulated cells were analysed as negative control (Unstim) and SEB as positive control. **C.** CCR6^+^ or **D.** CCR6^-^T-cells were stimulated with antigenic peptide pools derived from C. albicans (MP65) or Aspergillus (SHMT) and the co-expression of CD69 and the indicated cytokines analysed by flow cytometry. **E**: Co-expression of CD69 and IFN-γ in CCR6-Th-cells from AD-HIES patients (red bars, n=4-5) and healthy donors (HD, blue bars, n=8) stimulated with an EBV-derived peptide pool. Statistical significances between HD and HIES were calculated with unpaired t-tests.

<u>SUPPLEMENTARY FIGURE 6:</u>

Gating strategies for A: B-cell subsets and B: plamablasts. C: In 3 AD-HIES patients the B helper assay was first performed with total CCR6^+^T-cells and later repeated with CXCR3^-^CCR6^+^T-cells. IgG levels induced by total CCR6^+^Th-cells and by CXCR3^-^CCR6^+^T-cells from the same patients are connected by a line.

<u>SUPPLEMENTARY TABLE 1: LIST OF ANTIBODIES</u>

SUPPLEMENTARY TABLE 2: LIST OF ANTIGENS

## Supplementary METHODS

### Immunofluorecscence

Cells were fixed in 4% paraformaldehyde (PFA) for 10 min at room temperature, washed with PBS and blocking buffer (10% Fetal Calf Serum in PBS) was added for 1h at room temperature. Cells were then incubated with an anti-human CCR6 mouse antibody (1:50, R&D Systems) and an anti-CD4 rabbit monoclonal antibody (1:30, Abcam) overnight at +4°C. Subsequently, cells were washed, and incubated with a chicken anti-mouse IgG Alexa Fluor™ 568 (1:1000, ThermoFisher Scientific) and a goat anti-rabbit IgG Alexa Fluor™ 488 (1: 1000, ThermoFisher Scientific) for 1 hour at room temperature. Cells were washed with PBS, permeabilized with 0.1% Triton X-100 for 15 minutes, and incubated overnight with a rat anti-human IL-10 antibody (1:100, ThermoFisher, U.S.A). Cells were washed with PBS and incubated with a donkey anti-rat IgG Alexa Fluor™ 647 secondary antibody for 1 hour at room temperature. After washing with PBS, cells were counterstained with DAPI (Molecular Probes, ThermoFisher Scientific) and mounted using ProLong Diamond and ProLong Glass mounting reagents (ThermoFisher Scientific). Labelling control (no secondary antibody) was performed before starting the experiments. The isotype control was included in each experiment.

## Notes

### Competing Interest Statement

The authors have declared no competing interest.

### Summary of Updates

This version of the manuscript is more concise to enhance the clarity of the presented data

## REFERENCES

1. Sallusto F, Geginat J, Lanzavecchia A. Central memory and effector memory T cell subsets: function, generation, and maintenance. Annu Rev Immunol. 2004;22:745–763.

2. Rivino L, Messi M, Jarrossay D, Lanzavecchia A, Sallusto F, Geginat J. Chemokine receptor expression identifies Pre-T helper (Th)1, Pre-Th2, and nonpolarized cells among human CD4+ central memory T cells. J Exp Med. 2004;200(6):725–735.

3. Crotty S. Follicular helper CD4 T cells (TFH). Annual review of immunology. 2011;29:621–663.

4. Korn T, Oukka M, Kuchroo V, Bettelli E. Th17 cells: effector T cells with inflammatory properties. Semin Immunol. 2007;19(6):362–371.

5. Annunziato F, Cosmi L, Liotta F, Maggi E, Romagnani S. The phenotype of human Th17 cells and their precursors, the cytokines that mediate their differentiation and the role of Th17 cells in inflammation. International immunology. 2008;20(11):1361–1368.

6. Codarri L, Gyulveszi G, Tosevski V, et al. RORgammat drives production of the cytokine GM-CSF in helper T cells, which is essential for the effector phase of autoimmune neuroinflammation. Nature immunology. 2011;12(6):560–567.

7. Geginat J, Paroni M, Facciotti F, et al. The CD4-centered universe of human T cell subsets. Semin Immunol. 2013;25(4):252–262.

8. Singh SP, Parween F, Edara N, et al. Human CCR6+ Th Cells Show Both an Extended Stable Gradient of Th17 Activity and Imprinted Plasticity. J Immunol. 2023;210(11):1700–1716.

9. Zielinski CE, Mele F, Aschenbrenner D, et al. Pathogen-induced human TH17 cells produce IFN-gamma or IL-10 and are regulated by IL-1beta. Nature. 2012;484(7395):514–518.

10. Peters A, Lee Y, Kuchroo VK. The many faces of Th17 cells. Current opinion in immunology. 2011;23(6):702–706.

11. Maggi L, Santarlasci V, Capone M, et al. CD161 is a marker of all human IL-17-producing T-cell subsets and is induced by RORC. European journal of immunology. 2010;40(8):2174–2181.

12. Paroni M, Leccese G, Ranzani V, et al. An intestinal Th17 subset is associated with inflammation in Crohn’s Disease and activated by adherent-invasive Escherichia coli (AIEC). J Crohns Colitis. 2023.

13. Acosta-Rodriguez EV, Rivino L, Geginat J, et al. Surface phenotype and antigenic specificity of human interleukin 17-producing T helper memory cells. Nat Immunol. 2007;8(6):639–646.

14. Paroni M, Maltese V, De Simone M, et al. Recognition of viral and self-antigens by TH1 and TH1/TH17 central memory cells in patients with multiple sclerosis reveals distinct roles in immune surveillance and relapses. J Allergy Clin Immunol. 2017.

15. Duhen T, Campbell DJ. IL-1beta promotes the differentiation of polyfunctional human CCR6+CXCR3+ Th1/17 cells that are specific for pathogenic and commensal microbes. Journal of immunology. 2014;193(1):120–129.

16. Duhen T, Geiger R, Jarrossay D, Lanzavecchia A, Sallusto F. Production of interleukin 22 but not interleukin 17 by a subset of human skin-homing memory T cells. Nat Immunol. 2009;10(8):857–863.

17. Trifari S, Kaplan CD, Tran EH, Crellin NK, Spits H. Identification of a human helper T cell population that has abundant production of interleukin 22 and is distinct from T(H)-17, T(H)1 and T(H)2 cells. Nature immunology. 2009;10(8):864–871.

18. Morita R, Schmitt N, Bentebibel SE, et al. Human blood CXCR5(+)CD4(+) T cells are counterparts of T follicular cells and contain specific subsets that differentially support antibody secretion. Immunity. 2011;34(1):108–121.

19. Zhou L, Ivanov, II, Spolski R, et al. IL-6 programs T(H)-17 cell differentiation by promoting sequential engagement of the IL-21 and IL-23 pathways. Nature immunology. 2007;8(9):967–974.

20. Kastirr I, Maglie S, Paroni M, et al. IL-21 Is a Central Memory T Cell-Associated Cytokine That Inhibits the Generation of Pathogenic Th1/17 Effector Cells. Journal of immunology. 2014.

21. Korn T, Bettelli E, Gao W, et al. IL-21 initiates an alternative pathway to induce proinflammatory T(H)17 cells. Nature. 2007;448(7152):484–487.

22. Asano T, Khourieh J, Zhang P, et al. Human STAT3 variants underlie autosomal dominant hyper-IgE syndrome by negative dominance. J Exp Med. 2021;218(8).

23. Minegishi Y, Saito M, Tsuchiya S, et al. Dominant-negative mutations in the DNA-binding domain of STAT3 cause hyper-IgE syndrome. Nature. 2007;448(7157):1058–1062.

24. Tsilifis C, Freeman AF, Gennery AR. STAT3 Hyper-IgE Syndrome-an Update and Unanswered Questions. J Clin Immunol. 2021;41(5):864–880.

25. Milner JD, Brenchley JM, Laurence A, et al. Impaired T(H)17 cell differentiation in subjects with autosomal dominant hyper-IgE syndrome. Nature. 2008;452(7188):773–776.

26. Puel A, Cypowyj S, Marodi L, Abel L, Picard C, Casanova JL. Inborn errors of human IL-17 immunity underlie chronic mucocutaneous candidiasis. Curr Opin Allergy Clin Immunol. 2012;12(6):616–622.

27. Rivino L, Gruarin P, Haringer B, et al. CCR6 is expressed on an IL-10-producing, autoreactive memory T cell population with context-dependent regulatory function. J Exp Med. 2010;207(3):565–577.

28. Facciotti F, Larghi P, Bosotti R, et al. Evidence for a pathogenic role of extrafollicular, IL-10-producing CCR6(+)B helper T cells in systemic lupus erythematosus. Proc Natl Acad Sci U S A. 2020;117(13):7305–7316.

29. Carrabba M, Dellepiane RM, Cortesi M, et al. Long term longitudinal follow-up of an AD-HIES cohort: the impact of early diagnosis and enrollment to IPINet centers on the natural history of Job’s syndrome. Allergy Asthma Clin Immunol. 2023;19(1):32.

30. Kastirr I, Crosti M, Maglie S, et al. Signal Strength and Metabolic Requirements Control Cytokine-Induced Th17 Differentiation of Uncommitted Human T Cells. Journal of immunology. 2015;195(8):3617–3627.

31. Moschetti G, Vasco C, Clemente F, et al. Deep Phenotyping of T-Cells Derived From the Aneurysm Wall in a Pediatric Case of Subarachnoid Hemorrhage. Front Immunol. 2022;13:866558.

32. Romagnani S, Maggi E, Liotta F, Cosmi L, Annunziato F. Properties and origin of human Th17 cells. Molecular immunology. 2009;47(1):3–7.

33. Rutitzky LI, Bazzone L, Shainheit MG, Joyce-Shaikh B, Cua DJ, Stadecker MJ. IL-23 is required for the development of severe egg-induced immunopathology in schistosomiasis and for lesional expression of IL-17. J Immunol. 2008;180(4):2486–2495.

34. Gattinoni L, Ji Y, Restifo NP. Wnt/beta-catenin signaling in T-cell immunity and cancer immunotherapy. Clin Cancer Res. 2010;16(19):4695–4701.

35. Annunziato F, Cosmi L, Santarlasci V, et al. Phenotypic and functional features of human Th17 cells. J Exp Med. 2007;204(8):1849–1861.

36. Ma CS, Chew GY, Simpson N, et al. Deficiency of Th17 cells in hyper IgE syndrome due to mutations in STAT3. J Exp Med. 2008;205(7):1551–1557.

37. Geginat J, Vasco C, Gruarin P, et al. Eomesodermin-expressing type 1 regulatory (EOMES(+) Tr1)-like T cells: Basic biology and role in immune-mediated diseases. Eur J Immunol. 2023:e2149775.

38. Brockmann L, Soukou S, Steglich B, et al. Molecular and functional heterogeneity of IL-10-producing CD4(+) T cells. Nat Commun. 2018;9(1):5457.

39. van de Veen W, Kratz CE, McKenzie CI, et al. Impaired memory B-cell development and antibody maturation with a skewing toward IgE in patients with STAT3 hyper-IgE syndrome. Allergy. 2019;74(12):2394–2405.

40. Locci M, Havenar-Daughton C, Landais E, et al. Human Circulating PD-1CXCR3CXCR5 Memory Tfh Cells Are Highly Functional and Correlate with Broadly Neutralizing HIV Antibody Responses. Immunity. 2013.

41. Geginat J, Paroni M, Maglie S, et al. Plasticity of human CD4 T cell subsets. Front Immunol. 2014;5:630.

42. Okada S, Markle JG, Deenick EK, et al. IMMUNODEFICIENCIES. Impairment of immunity to Candida and Mycobacterium in humans with bi-allelic RORC mutations. Science. 2015;349(6248):606-613.

43. Steward-Tharp SM, Laurence A, Kanno Y, et al. A mouse model of HIES reveals pro-and anti-inflammatory functions of STAT3. Blood. 2014;123(19):2978–2987.

44. McGeachy MJ, Bak-Jensen KS, Chen Y, et al. TGF-beta and IL-6 drive the production of IL-17 and IL-10 by T cells and restrain T(H)-17 cell-mediated pathology. Nat Immunol. 2007;8(12):1390–1397.

45. Yang L, Anderson DE, Baecher-Allan C, et al. IL-21 and TGF-beta are required for differentiation of human T(H)17 cells. Nature. 2008;454(7202):350–352.

46. Manel N, Unutmaz D, Littman DR. The differentiation of human T(H)-17 cells requires transforming growth factor-beta and induction of the nuclear receptor RORgammat. Nature immunology. 2008;9(6):641–649.

47. Steinfelder S, Floess S, Engelbert D, et al. Epigenetic modification of the human CCR6 gene is associated with stable CCR6 expression in T cells. Blood. 2011;117(10):2839–2846.

48. Sallusto F, Lenig D, Mackay CR, Lanzavecchia A. Flexible programs of chemokine receptor expression on human polarized T helper 1 and 2 lymphocytes. J Exp Med. 1998;187:875–883.

49. Freeman AF, Holland SM. Clinical manifestations, etiology, and pathogenesis of the hyper-IgE syndromes. Pediatr Res. 2009;65(5 Pt 2):32R–37R.

50. Xu H, Agalioti T, Zhao J, et al. The induction and function of the anti-inflammatory fate of TH17 cells. Nat Commun. 2020;11(1):3334.

51. Ma CS, Wong N, Rao G, et al. Unique and shared signaling pathways cooperate to regulate the differentiation of human CD4+ T cells into distinct effector subsets. J Exp Med. 2016;213(8):1589–1608.

52. Sabat R, Grutz G, Warszawska K, et al. Biology of interleukin-10. Cytokine Growth Factor Rev. 2010;21(5):331–344.

