## Supplementary figures and images for "AD-HIES patients retain preTh17-cells that produce IL10 with opportunistic pathogens and induce IgE"

### Supplementary figures

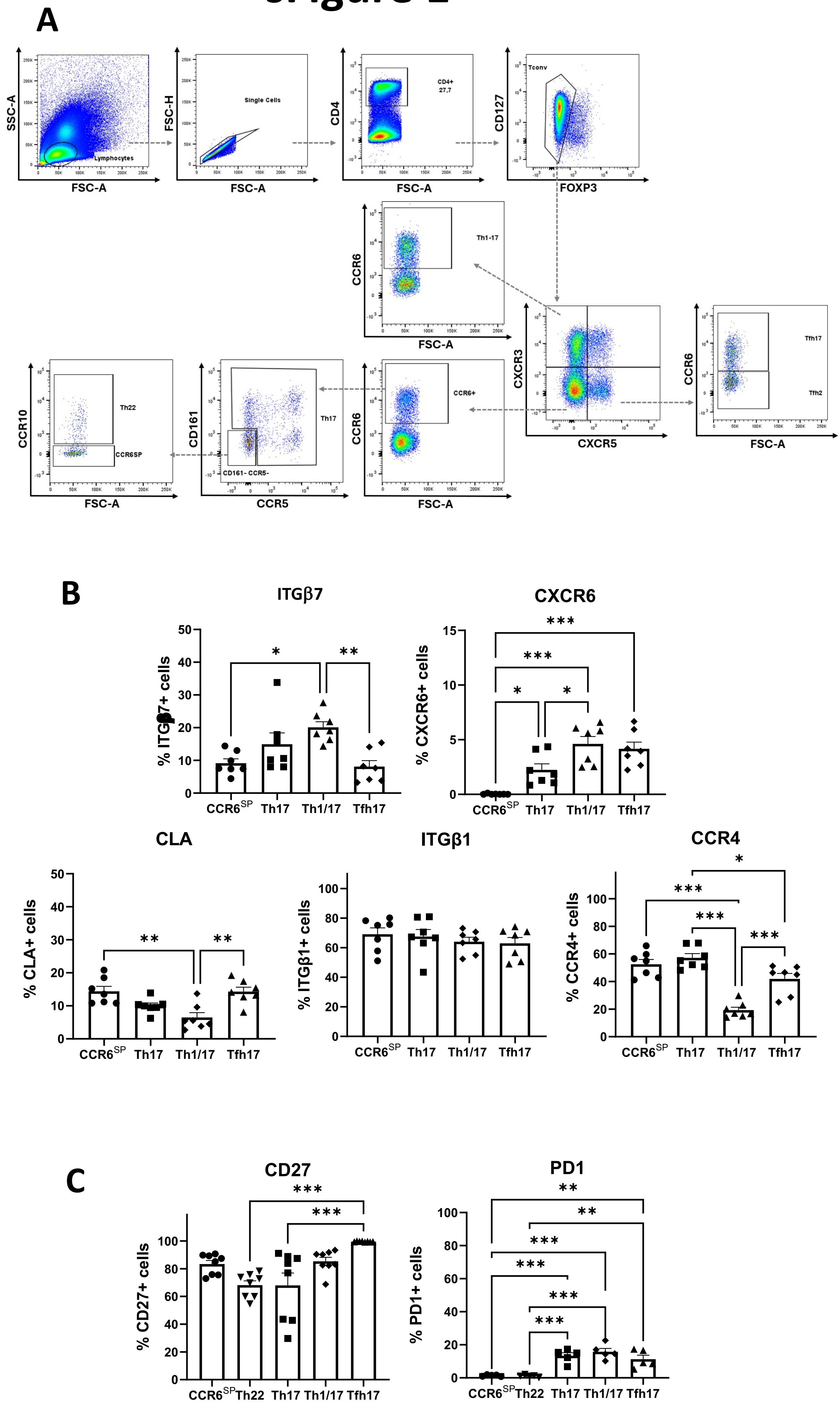

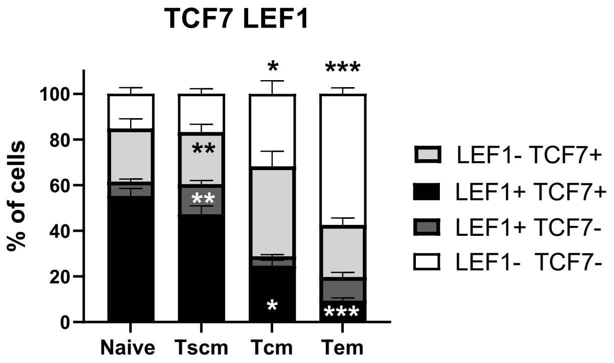

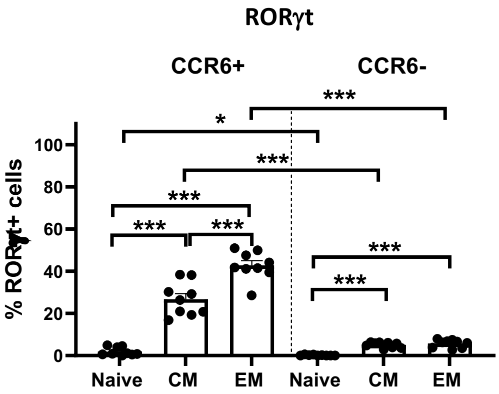

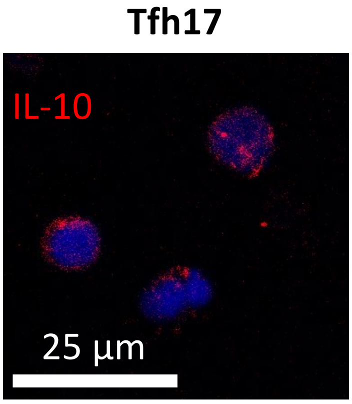

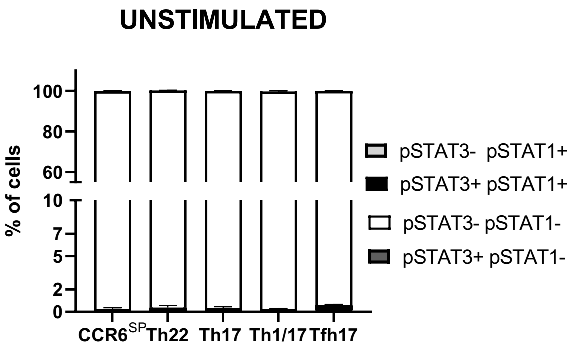

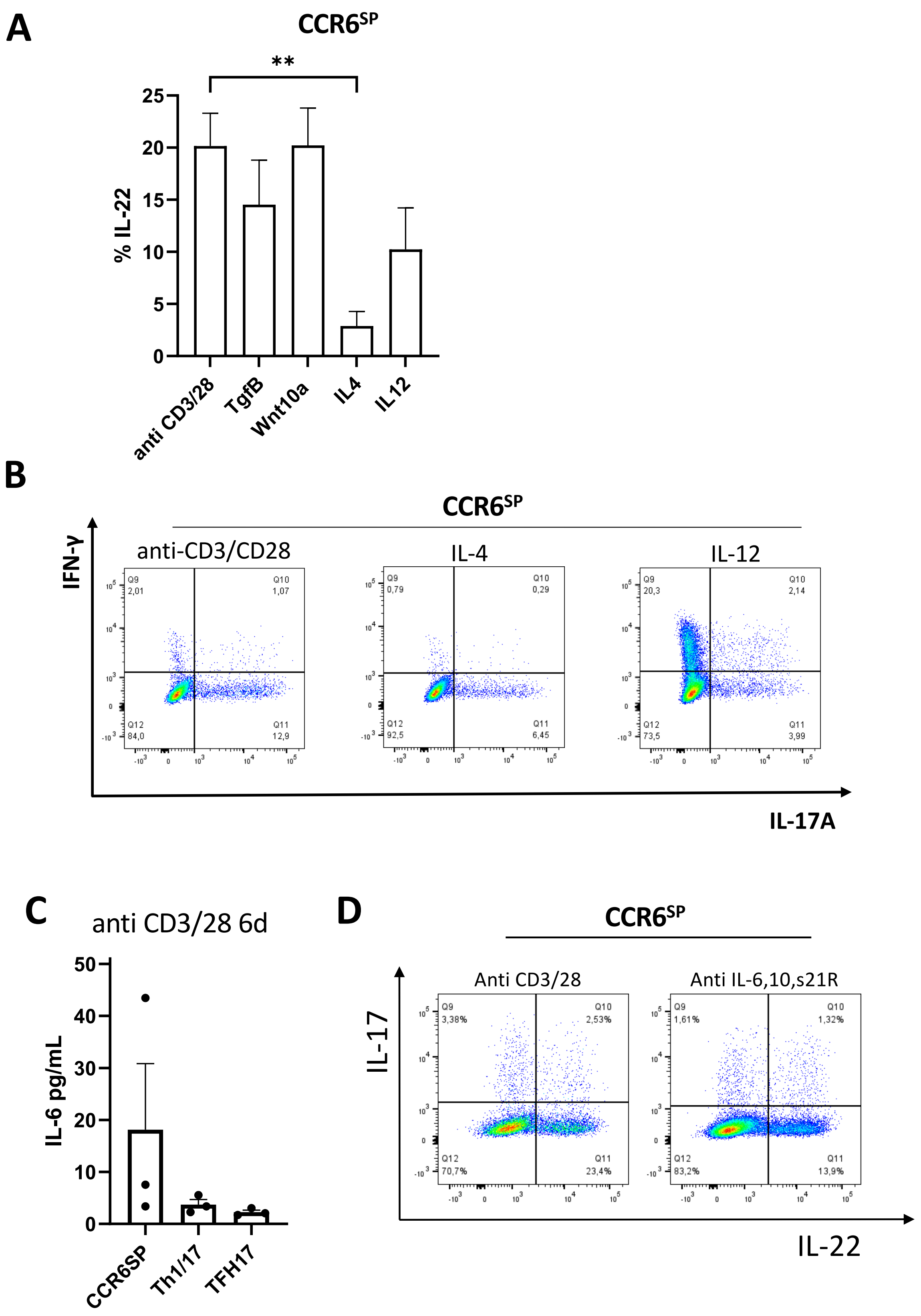

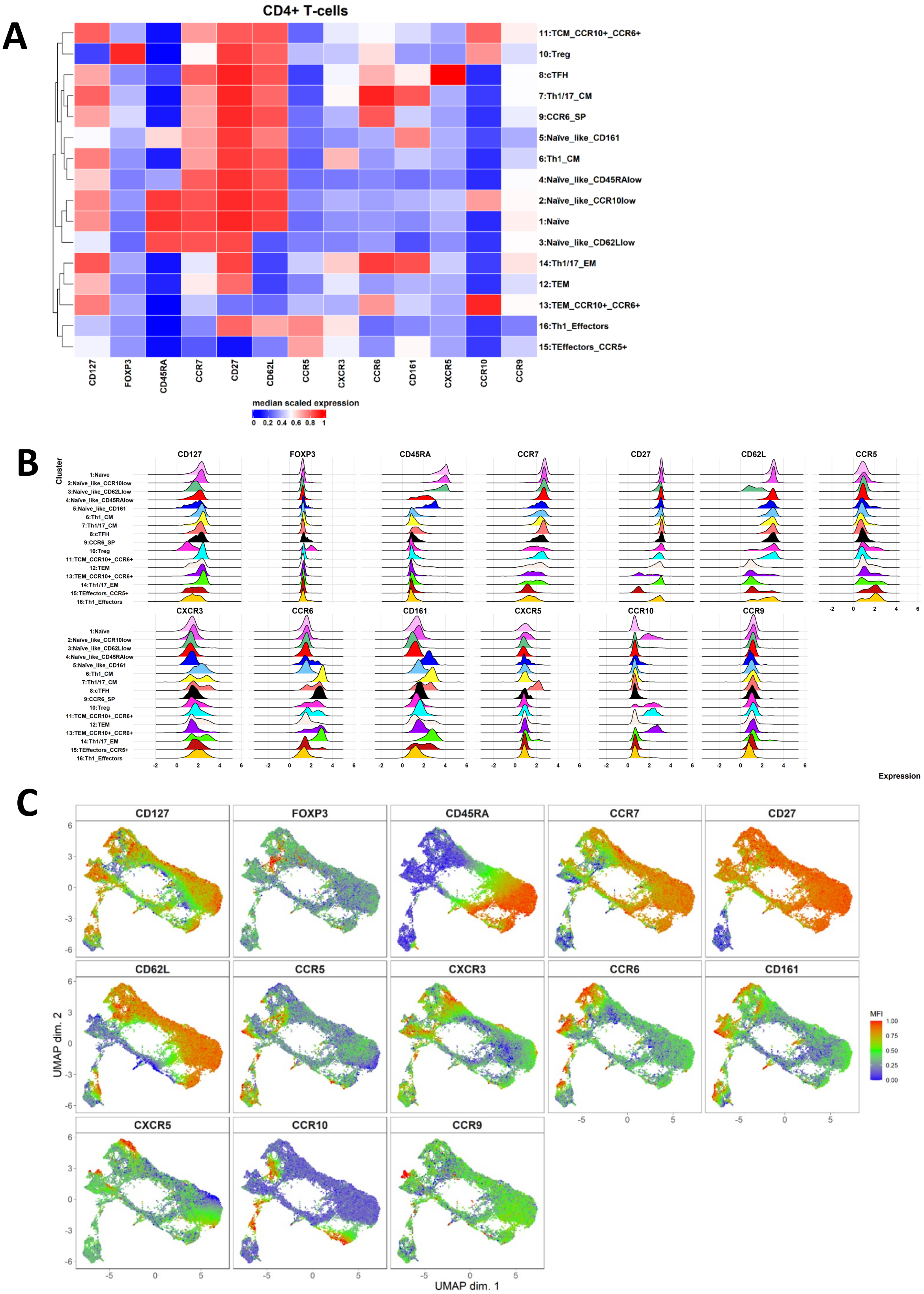

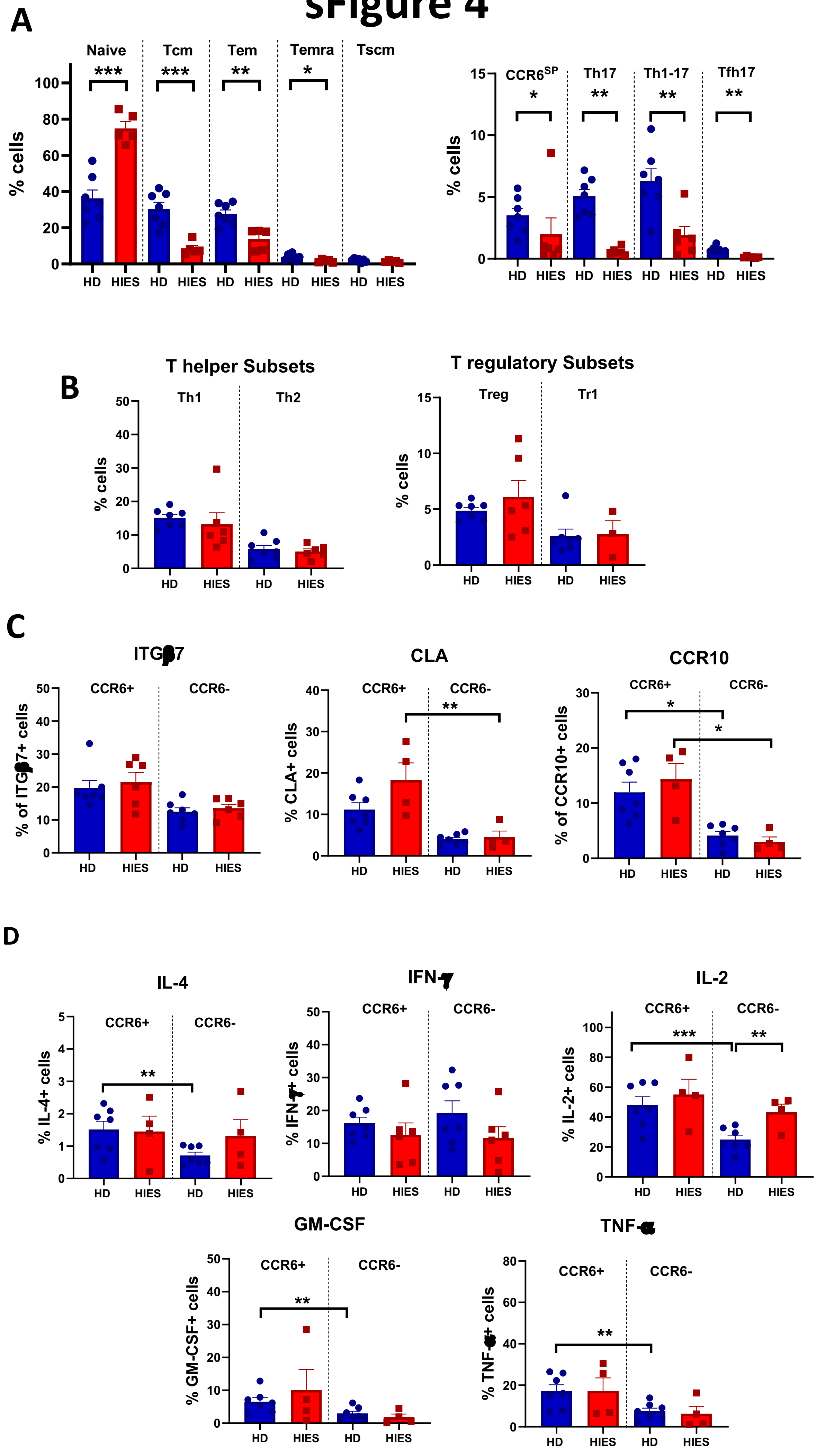

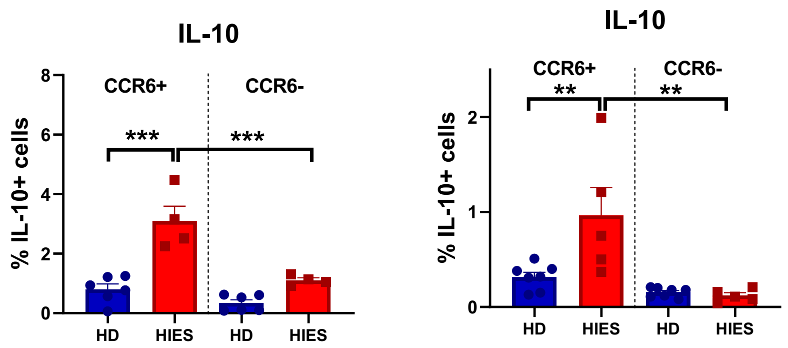

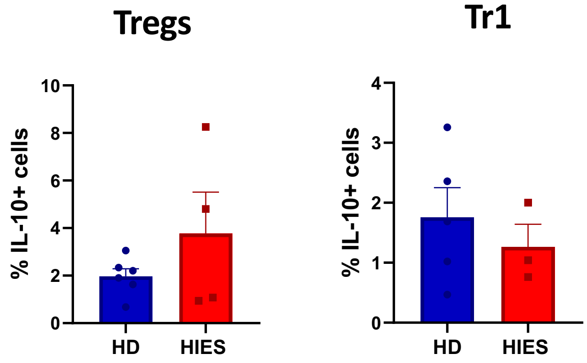

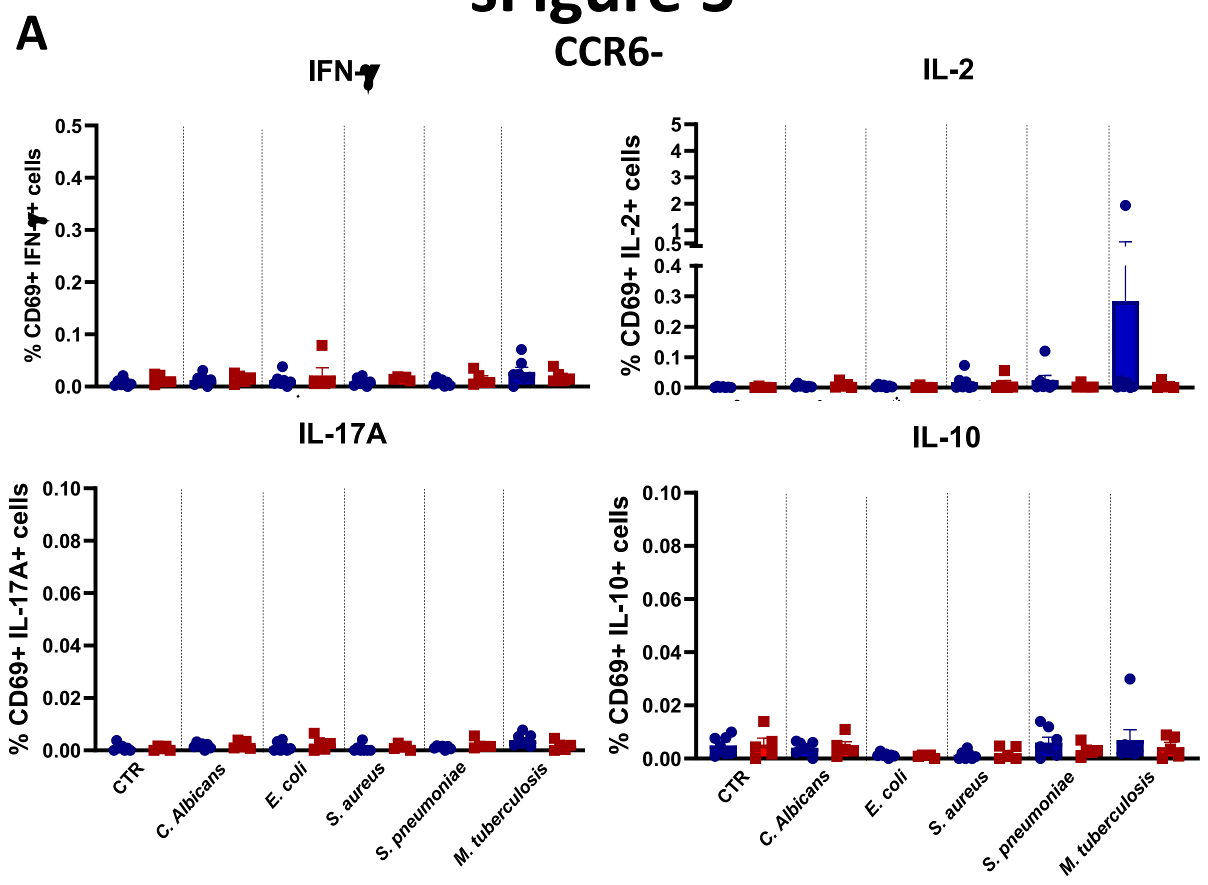

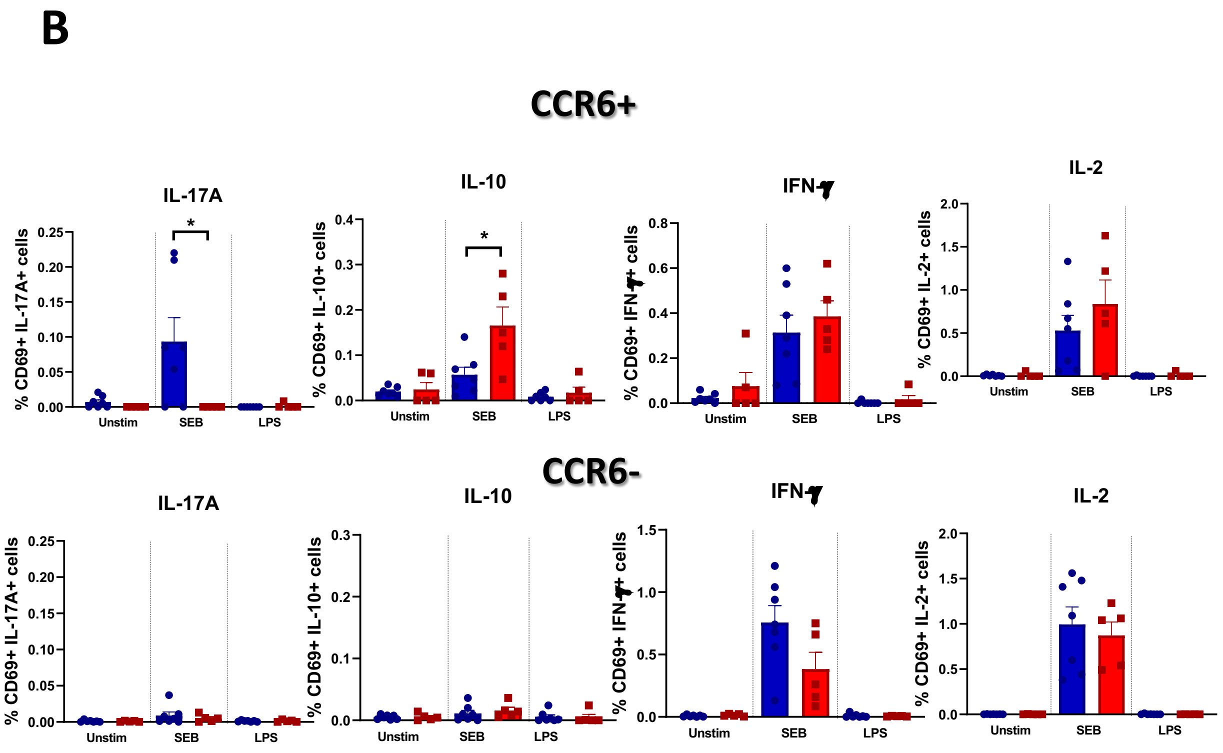

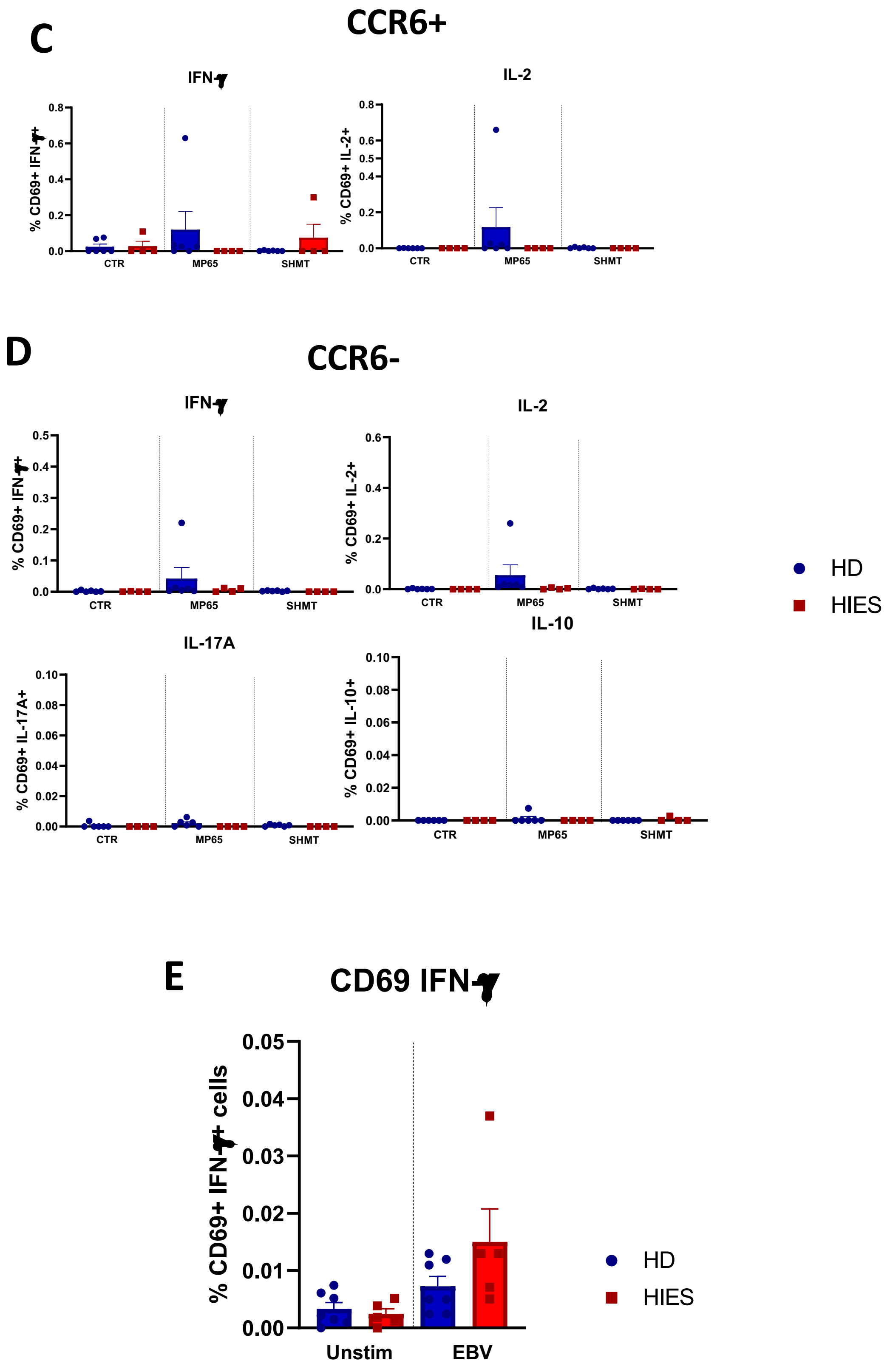

#### sFigure 6

**A**

B cells

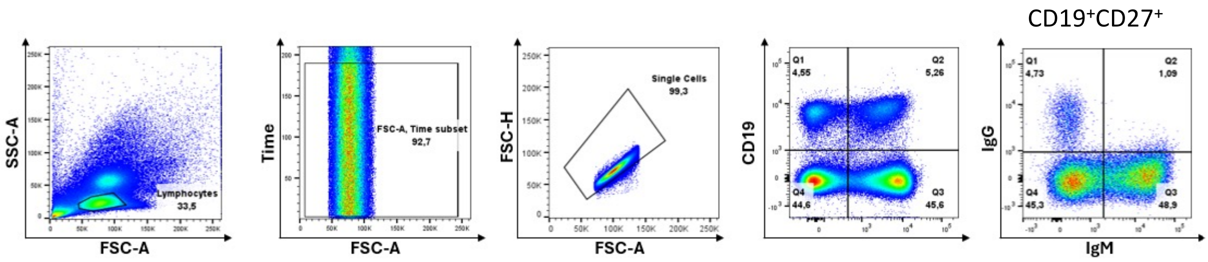


**B**

Plasmablasts

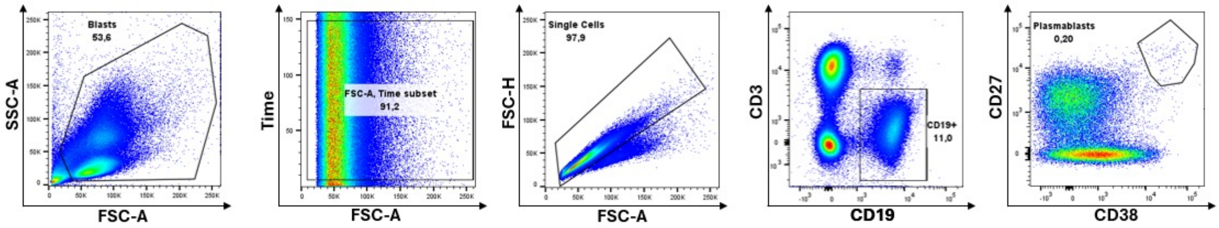


**C**

Th + SEB

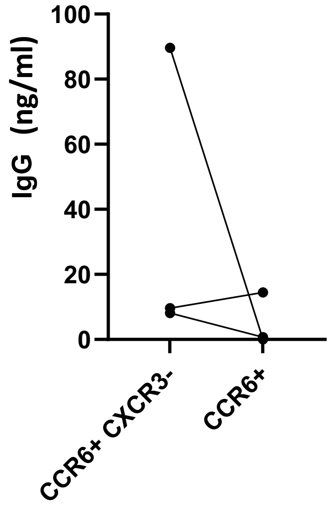
